# Autopolyploidization presents a transient and potential-rich window of increased transcriptional plasticity in *Arabidopsis arenosa*

**DOI:** 10.64898/2026.06.16.732565

**Authors:** Sonia Celestini, Eliška Trávníčková, Marek Brindzák, Filip Kolář

## Abstract

Whole-genome duplication (WGD, polyploidization) is a pervasive feature of Eukaryote evolution and often viewed as a source of evolutionary success and novelty, meaning a macromutation leading to higher fitness (i.e. “hopeful monsters”). Yet, the mechanisms behind the (occasional) success of nascent polyploids remain still elusive, especially from a transcriptomic point of view. Theory suggests that duplicated genetic networks are characterised by enhanced redundancy and higher output variation, promoting the exploration of the adaptive landscape during stressful times. Artificially synthesized neo-polyploid mutants provide an exciting system to test this, however, empirical studies comparing co-expression network patterns between natural and synthetic ploidies of the same species in an evolutionary context are lacking. Here we compare diploid, synthetic and naturally established autotetraploid populations of *Arabidopsis arenosa* to investigate short- versus long-term effects of polyploidy on gene expression complexity and plasticity under water deficiency stress. Transcriptomic profiling revealed that synthetic neo-tetraploids explored the broadest expression space and exhibited the highest number of stress-responsive genes. Co-expression network analyses demonstrated that the network of neo-tetraploids was fragmented into multiple highly connected modules, with stress-responsive genes preferentially acting as inter-modular “bridges”. In contrast, diploids and established tetraploids exhibited lower expression variation, more modular architectures, with stress response genes embedded within well-defined modules. Moreover, synthetic tetraploids displayed the highest number of modules correlated with plant fitness proxy suggesting higher output variance resulting from the transcriptional shock. Together, our results indicate that WGD induces a transient phase of transcriptomic expansion and network disorganization that broadens the phenotypic landscape, followed by evolutionary stabilization and finally retention of some advantageous novelties in established polyploids. This supports the view of neo-polyploids as “hopeful monsters”, in which short-term instability creates a window of enhanced variability and plasticity with long-term evolutionary potential.

**Significance:** Since the early concept of polyploids as “hopeful monsters,” biologists have hypothesized that whole-genome duplication can generate novel phenotypes and facilitate adaptation to environmental challenges. Yet the mechanisms linking genome doubling to evolutionary innovation remain poorly understood. By comparing diploid, synthetic autotetraploid, and naturally established autotetraploid populations of Arabidopsis arenosa, we show that newly formed polyploids undergo a transient phase of expanded transcriptomic variation and extensive regulatory network rewiring. In contrast, established polyploids exhibit a more stable and modular network architecture. Our results provide empirical support for a long-standing evolutionary hypothesis, showing how genome duplication can temporarily broaden the range of possible phenotypes before subsequent stabilization through evolution.

## Introduction

Within species whole-genome duplication (WGD), or autopolyploidy—here also called polyploidy—is suggested to play a central role in plant evolution, shaping lineage diversification, speciation and adaptation (Alix et al. 2017; Van de Peer et al. 2017; Fox et al. 2020). For example, polyploids frequently exhibit novel or enhanced phenotypes, including increased vigour or cellular dimension, altered physiology, and improved stress tolerance (Bomblies 2020; Van de Peer et al. 2020; Duan et al. 2023). So far, mostly the duplication of genes *per se* has been associated with the success of polyploids, thanks for example to immediate heterosis or long-term genetic robustness and functional divergence of each copy (Otto and Whitton 2000; Soltis and Soltis 2000; Comai 2005; Parisod et al. 2010; De Smet and Van de Peer 2012). Alternative scenarios, such as rewiring of the cellular networks following WGD, may also underpin evolutionary potential of nascent autopolyploids; however, empirical evidence is lacking (De Smet and Van de Peer 2012; Parisod 2024).

Recent work mostly focused on computational frameworks aimed at mimicking the biological evolution of different ploidy cytotypes. Studies using digital organisms (Yao et al. 2019), virtual cells (Cuypers and Hogeweg 2014) and artificial ancestral gene regulatory networks (aGRNs) (Ebadi et al. 2023), synergistically demonstrated a context-dependent contribution of WGD to the evolutionary success of a nascent polyploid lineage: non-duplicated networks perform better in stable environments, while duplicated networks show an advantage under environmental stress or turmoil. Mechanistically, WGD doubles the number of network nodes (e.g., genes, alleles, proteins) and exponentially increases the number of edges (interactions), enhancing redundancy, complexity, and possibly modularity (Fusco et al. 2010; Doyle and Coate 2019). Despite probably initially maladaptive, due to so-called “genomic shock” (Doyle and Coate 2019; Scarrow et al. 2021), these properties are suggested to expand the range of possible phenotypic responses and promote evolutionary potential by enabling exploration of novel adaptive peaks (Yao et al. 2019; Ebadi et al. 2023; Parisod 2024). Furthermore, the expansion of gene regulatory networks through duplication introduces functional redundancy, which buffers against deleterious variants and preserves core ancestral functions while facilitating the emergence of novel regulatory connections—an architecture particularly advantageous in variable or stressful environments (Cuypers and Hogeweg 2014).

Experimental evidence in *Saccharomyces cerevisiae* shows that polyploid strains adapted more rapidly than diploids under selective pressure, however Selmecki et al. (2015) and Scott et al. (2017) showed that this was due to enhanced access to beneficial mutations and increased genomic flexibility long-term, while the relatively more short-term role of network rewiring is still unexplored. A recent comparative genomic work has demonstrated that ancient WGD events played an major role in shaping the topology of GRNs in angiosperms (Almeida-Silva and Van de Peer 2023). Here, polyploidy likely conferred a short-term selective advantage by increasing the frequency of network motifs (simple genetic circuits that perform elementary regulatory functions) and therefore putatively increasing robustness (i.e., greater number of nodes, edges, and interactions thereof) in GRNs, providing raw genetic material (i.e., network nodes) for evolving novel regulatory interactions (Almeida-Silva and Van de Peer 2023). Alternatively, the greater number of motifs amplified their combinatorial effect and therefore output variation, which allowed greater jumps in the fitness landscape as suggested by simulations (Ebadi et al. 2023). In addition, they observed that these motifs tend to get lost over time, probably because duplicated genes are simply lost for fractionation or because networks are rewired.

Recent post-WGD evolution of gene (co-)expression in general, and the regulatory mechanisms underlying ploidy-specific responses to environmental perturbations in particular, remain incompletely understood, in spite of the ubiquity of plant systems encompassing diploids and their recent intraspecific polyploid derivatives (Rice et al. 2015; Kolář et al. 2017). Comparative studies between diploids and their synthetic autopolyploid relatives were mostly based on comparison of differential expression patterns, and often demonstrated little to moderate effects of WGD on gene expression patterns — for example in *Medicago sativa* (Rosellini et al. 2016; Santoro, Marconi, et al. 2025; Santoro, Anderson, et al. 2025), in *Capsella* (Duan et al. 2023), in potato (Stupar et al. 2007; Aversano et al. 2015), *Citrus limonia* (Allario et al. 2011), *Chrysanthemum lavandulifolium* (Gao et al. 2016), and *Arabidopsis thaliana* (Yu et al. 2010; Liu et al. 2017). The highly variable outcomes of genome duplication at the transcriptional level may reflect genotype-specific effects or stochastic consequences of the genomic shock that accompanies polyploidization, which may require extended time for adjustment and adaptation to the altered cellular context (Vyas et al. 2007; Yu et al. 2010; Doyle and Coate 2019). Indeed, even if neo-polyploids do not usually display large-scale transcriptional changes compared to their diploid progenitors, established polyploids may nonetheless retain and exploit novel transcriptional variation introduced by WGD, which could underlie their frequently reported greater capacity to cope with environmental stresses (Van de Peer et al. 2020; Tossi et al. 2022). Studies comparing gene expression between established natural polyploids and their close diploid relatives, especially under stress conditions, are, however, rare and discordant (Church and Spaulding 2009; Zhou et al. 2019; Srikant et al. 2024). Moreover, interploidy comparisons of expression patterns specifically related to stress responses are limited to cultivars or model species (Chao et al. 2013; Yang et al. 2014; Wang, Fan, et al. 2022; Wang, Wang, et al. 2022), leaving unknown the effect of post-WGD evolution in natural context. In summary, comparisons between diploids and synthetic newly formed polyploids often leave unclear whether the observed changes are adaptive or maladaptive and whether they will persist as evolution proceeds. Conversely, when contrasting diploids with naturally established polyploids, it becomes difficult to disentangle which differences stem directly from genome duplication versus which arose through independent post-WGD evolutionary processes during cytotype divergence. We, however, miss studies providing in-depth comparison of gene (co-)expression between all three categories from an ecological and evolutionary point of view, leaving unclear the effect of WGD *per se* vs. post-WGD differentiation on gene expression evolution.

*Arabidopsis arenosa* provides an ideal system to investigate the consequences of polyploidy in nature, as this herbaceous plant species occurs both as diploid and naturally established autotetraploid populations across Europe (Kolář et al. 2016; Monnahan et al. 2019; Padilla-García et al. 2023). Notably, the autotetraploid lineage of *A. arenosa* is thought to have originated ∼19,000–31,000 generations ago from a single diploid ancestor lineage in the Western Carpathians, followed by an extensive postglacial expansion (Monnahan et al. 2019) where both ploidies adapted to diverse set of abiotic natural challenges along broad temperature and precipitation gradients (high latitude and high alpine habitats; Knotek et al. 2020; Wos et al. 2023; Bohutínská et al. 2024) as well-as local substrate-driven challenges (drought-prone substrates; Guggisberg et al. 2018; Preite et al. 2019; Konečná et al. 2021). Importantly, both ploidies still co-exist in the area of putative WGD origin in the Western Carpathians providing an exciting opportunity to compare diploids with their established polyploid outcomes occupying the same environmental niche (Morgan et al. 2020). Recent work in *A. arenosa* has shown that WGD can rapidly alter chromatin accessibility and differential expression patterns in synthetic autopolyploids, however, in natural tetraploids most expression patterns, as probably previously detrimental, reverted towards a diploid-like state (Srikant et al. 2024). However, as the natural tetraploid population TBG came from evolutionarily derived, allopatric lineage adapted to very specific and challenging habitat (railway tracks) it is unclear to which extent a lineage-specific post-polyploid evolution contributed to such a pattern. These results thus provided intriguing mechanistic explanation partly underlying the altered expression patterns, yet we miss an evolutionary and ecological perspective on the adaptive potential of this altered expression profile under varying levels of environmental stress.

In this study we explore this issue by comparing gene (co-)expression in leaves of diploids, synthetic neo-tetraploids, and established autotetraploids of *A. arenosa*, representing a single evolutionary lineage comprising genetically and ecologically close populations from a natural contact zone in Western Carpathians. All cytotypes were subjected to limited water availability stress – a globally relevant factor which *A. arenosa* readily faces across its native range (Morgan et al. 2020; Padilla-García et al. 2023). We identify drought-responsive genes (differentially expressed genes—DEGs) in each ploidy group and perform weighted gene co-expression network analysis (WGCNA) to identify co-expressed gene clusters (modules) and quantify connectivity measures informative on the network properties. These metrics allow insights into how stress-responsive genes (DEGs) are embedded within or between transcriptional modules, providing a systems-level perspective on gene regulatory architecture and plasticity.

Specifically, we ask: (i) How do differential expression patterns and topology of co-expression networks differ across diploid, neo-, and established tetraploid *A. arenosa* plants? (ii) How are drought-responsive genes (DEGs) integrated within these networks in each ploidy? (iii) Does neo-polyploidy imply novel, transgressive expression phenotypes conferring increased transcriptional disparity and plasticity as compared to the other ploidies? (iv) Are any of these changes preserved in the established natural tetraploids? By combining experimental manipulations of synthetic polyploids and natural populations with high-resolution co-expression network analysis, this study provides novel insights into how genome duplication influences the modularity and flexibility of stress responses both in short and long-term, with a broader aim to understand the mechanisms underlying evolutionary success of polyploid populations we observe in the wild.

## Material and Methods

### Dataset and experimental treatment

To enable a meaningful comparison between diploid and established tetraploid samples, we focus on genetically and ecologically close natural diploid and autotetraploid populations of *Arabidopsis arenosa* from the Western Carpathians. In this area autotetraploids singularly originated from a diploid ancestor approximately 20-30 k generations ago (Monnahan et al. 2019), and both ploidies still co-occur here since then, sharing similar ecological niche (Wos et al. 2019; Morgan et al. 2020; Padilla-García et al. 2023). Specifically, we collected seeds from two geographically close populations (26 km) differing in ploidy, living at a similar altitude and environment in central Slovakia. Diploid individuals (2n = 2x = 16) were sampled from the population AA012 in Špania Dolina (48.80625 N, 19.13147222 E), while established natural tetraploids (2n = 4x = 32) individuals were collected from the nearby population AA225 south of Ružomberok (49.017836 N, 19.283829 E). In this way, we aim to ensure that differences observed between diploid and tetraploid individuals are primarily attributable to ploidy level status rather than to a broader genomic or ecological divergence. We collected seeds from ∼15 mother plants per each population and stored them in microtubes, afterwards we stratified them at 4°C for several weeks.

To minimize maternal effects of the original sampling sites, we first grew one (F1) generation from the locally collected seeds under controlled conditions. The plants were cultivated at all stages in a walk-in growth chamber (FYTOSCOPE FS-WI-GU, PSI: Photon Systems Instruments, Ltd.) under the following conditions: day – 16 hours at 20°C; night – 8 hours at 15°C; relative humidity maintained at approximately 65–70%; and light intensity set to ∼145 μmol⋅m=²⋅s=¹. After stratification, ∼50 seeds per mother were sown on Petri dishes lined with moist filter paper and misted with water every 2–3 days. Seedlings that developed both cotyledons and roots were transplanted 14 days after germination into larger pots (7 × 7 × 8 cm). After 10 weeks, the plants were vernalized for 6 weeks (conditions: day – 8 hours at 6°C; night – 16 hours at 6°C) and then manually crossed with an individual from different seed family from the same population. In this way, about 12 new families representatively capturing the diversity of each natural ploidy source population were created.

The resulting F2 seeds were cold-stratified and directly sown into multipot trays filled with soil (2.5 × 2.5 × 3 cm cells). Seedlings were watered from below three times per week. After 45 days, young rosettes were transplanted into larger pots (7 × 7 × 8 cm), each filled with 120 g of soil (sterilized mix of peat, sand, and perlite; Forestina substrate for sowing). At day 60 after sowing, plants were divided into two treatment groups in a way that progeny of each F1 family was represented in each treatment by at least one individual, but the assignment of individuals to these treatment groups was random. The first group received the control treatment, where soil moisture was kept at approximately 50% of field capacity through regular watering with 50 g of water every 3-5 days. The second group was exposed to the water limitation (drought) treatment, meaning they were watered with 50 g only upon visible signs of leaf wilting to ensure consistent drought stress (as done in Bouzid et al. 2019). We focused on water limitation as it is a ubiquitous natural stressor that can be reliably experimentally manipulated, it is broadly relevant under ongoing global change in both natural and agricultural systems, including also *A. arenosa* whose populations of both ploidy levels occupy broad gradient of precipitation (Morgan et al. 2020; Padilla-García et al. 2023).

For each cultivated plant, we scored the total number of flowers produced until the end of the experiment (∼200^th^ day of cultivation), and the rosettes diameter at the time of RNA collection (100^th^ day). These phenotypic data were then statistically analysed in R v.4.4.2. We tested the effect of ploidy and water treatment, and their interaction, on the phenotypes using linear models with the function lm() from the package lme4 1.1-37 (Bates et al. 2015). The input data was normalised using the package bestNormalize v.1.9.1 (Peterson and Cavanaugh 2020) to meet model assumptions. Normality of residuals and homogeneity of variance were visually inspected using diagnostic plots.

### Polyploid synthesis

Because synthetic neo-polyploid individuals share an almost identical genetic background with their diploid progenitors, comparisons between these cytotypes allow for a direct assessment of the immediate effects of whole-genome duplication, independent of evolutionary divergence. In contrast, comparisons involving established natural polyploid populations reflect both the legacy of genome duplication and the subsequent evolutionary changes, such as allele substitutions, gene silencing, and evolved regulatory rewiring. To disentangle these processes, we generated synthetic autotetraploids from natural diploid individuals from the same population (AA012) via oryzalin, a mitotic poison commonly used as herbicide of the dinitroaniline class. About 150 seeds belonging to the F1 generation (field-collected) were sown and germinated as described above. Then, seedlings aged between 10 and 14 days, at the stage when a small tip with meristematic tissue becomes visible between the cotyledons, were treated with 40 µL of a 500 µM oryzalin solution prepared in 5% dimethyl sulfoxide (DMSO). After 48 hours, the seedlings were thoroughly washed with water. A subset of the treated plants was then transferred into multipot trays and grown under the same conditions described in the previous section to generate a non-treated F2 generation that was then used for our experiment. To assess ploidy status of all oryzalin-treated plants, relative DNA content was estimated using a standard two-step nuclei isolation protocol with Otto buffers followed by flow cytometry, as described by Doležel et al. (2007). For each sample, leaf tissue was co-chopped with an internal standard (*Carex acutiformis*) of known genome size (Temsch et al. 2022) and stained with 4′,6-diamidino-2-phenylindole (DAPI). DNA ploidy levels were inferred from the relative ratio of the first fluorescence peak of the sample to that of the internal standard, which averaged 0.36 for diploid individuals and 0.73 for tetraploids. Samples were processed using a CytoFLEX Flow Cytometer (Beckman Coulter, Inc.) and were analysed with CytExpert software for the CytoFLEX platform (Beckman Coulter, Inc.). The diploid individuals from the F2 generation of the oryzalin-treated plants (so where the treatment did not lead to polyploidization) were used as a control of the polyploidization phenotypic effect (Supplementary fig. S1).

### Transcriptome sampling and processing

To representatively cover natural diversity within genetically highly variable outcrossing populations of *A. arenosa* (Vlček et al. 2025), we characterised transcriptomes of 30 individuals per each cytotype (15 per treatment). We collected the middle rosette leaf from each F2 plant at a consistent developmental stage (100 days after cultivation starts), i.e. when the rosette growth reached a plateau (Supplementary fig. S2). Immediately after collection, the leaf samples were snap-frozen in liquid nitrogen and stored at –80°C until RNA extraction. We extracted total RNA using the NucleoSpin RNA Plus kit (Macherey-Nagel), following the manufacturer’s protocol. To assess RNA quality, we measured purity and concentration with a NanoDrop One spectrophotometer (ThermoFisher Scientific) and evaluated RNA integrity using an Agilent 2100 Bioanalyzer (Agilent Technologies). We retained only those samples with RNA Integrity Number (RIN) scores above 7 for downstream processing. Library preparation and sequencing were performed by Macrogen Europe B.V. Libraries were prepared using the Illumina TruSeq Stranded mRNA Kit (Illumina) using 1 µg total of RNA per sample, and sequencing was carried out on the Illumina NovaSeq X platform, generating 150 bp paired-end reads.

We obtained a minimum of 15 million paired-end reads for each sample. We assessed the quality of raw sequencing reads using FastQC v.0.11.9 (Andrews 2010) and compiled summary reports with MultiQC v.1.14 (Ewels et al. 2016). Quality control included checks for per-base sequence quality, GC content, adapter contamination, and sequence duplication levels. We then mapped the reads to the diploid *Arabidopsis arenosa* reference genome from the same Western Carpathians lineage (Bramsiepe et al. 2023) using STAR v.2.7.10 (Dobin and Gingeras 2015). The genome index was generated using the STAR --runMode genomeGenerate function with --genomeSAindexNbases set to 12, --sjdbOverhang set to 149 and using the corresponding annotation file in GTF format. Paired-end reads were aligned using default parameter except for the following key options: --runThreadN 10, --outSAMtype BAM SortedByCoordinate, --quantMode GeneCounts, -seedSearchStartLmax 30, --outFilterScoreMinOverLread 0, --outFilterMatchNminOverLread 0. Alignment quality metrics such as mapping rate, mismatch rate, and number of multimapping reads were compiled across all samples using STAR log files. Based on this, one established tetraploid (4x) sample under drought conditions was excluded from the dataset due to very low RNA quality and mapping (Supplementary Table S1).

### Differential gene expression analysis

In order to detect drought stress responsive genes, differential gene expression analysis was performed using the DESeq2 v.1.38.3 package (Love et al. 2014) in R v.4.2.2. For each sample, raw gene-level read counts were obtained from STAR’s ReadsPerGene.out.tab file, using the fourth column, which corresponds to reverse stranded read counting. Genes with fewer than 10 counts in at least 50% of the samples were filtered out prior to analysis. We used the DESeqDataSetFromMatrix() function to create a DESeq2 object, incorporating the experimental design: ∼ ploidy + treatment + ploidy:treatment. Normalization and dispersion estimation were carried out using DESeq2’s default method, which uses a median-of-ratios approach to correct for differences in library size. Differential expression testing was conducted using Wald test and p-values were adjusted for multiple testing using the Benjamini–Hochberg method. Genes with an adjusted p-value (FDR) < 0.1 and a log_2_ fold change ≥ 1 (or ≤ –1) were considered significantly differentially expressed. To explore global patterns of gene expression and assess sample clustering, we performed principal component analysis (PCA) using variance-stabilized data. Raw count data were transformed using the variance-stabilizing transformation (VST) implemented in the vst() function from DESeq2, with the parameter blind=TRUE to avoid bias from experimental design. We used the plotPCA() function in DESeq2 and ggplot2 v.3.5.2 package (Wickham 2016) to generate PCA plots based on the top 500 most variable genes across all samples. Differences in global transcriptomic profiles between treatments and ploidies visualised by PCA were then tested using permutational multivariate analysis of variance (PERMANOVA) based on Euclidean distances. Significance was assessed using 999 permutations implemented in the adonis2 function of the vegan v.2.7-2 R package (Oksanen et al. 2025). Homogeneity of multivariate dispersions was verified using the betadisper test. Finally, volcano plots were generated for the same data using the package EnhancedVolcano v.1.16.0 (Blighe et al. 2022).

### Multivariate disparity analysis

To quantify and compare the extent of phenotypic space (transcriptomic variation) occupied by each ploidy group in each treatment, we calculated multivariate disparity using the dispRity v.1.9 R package (Guillerme 2018). Principal component analysis (PCA) was first performed on the variance-stabilized expression data produced as described above, and the first 15 principal components (PCs), representing the majority (55%) of the variance in the dataset, were retained for disparity estimation. Disparity was then calculated for each ploidy-treatment group as the sum of variances (overall spread in multivariate space) bootstrapping each subset 1,000 times using the “full” algorithm. The resulting bootstrap distributions of disparity values for each group were plotted using the default plot() function from dispRity. Additionally, to visualize the distribution of transcriptomic profiles in reduced dimensional space, the first two PCs were plotted using 2D kernel density contours in ggplot2 for each combination of ploidy and treatment.

### Plasticity analysis

Further, we aimed at directly quantifying a possible difference in plasticity between ploidies. In order to do so, we scored plasticity at the level of the transcriptomic phenotype as the magnitude of environmentally induced change in global gene expression profile of a genotype. We indeed measured the multivariate displacement in gene expression profiles between pairs of full sibling plants (coming exactly from the same parents) grown under contrasting water regimes. We focused on diploid (15 pairs) and synthetic tetraploid (19 pairs) plants in order to asses net effects of ploidy differences on transcription plasticity. As input we used the first 15 principal components calculated from the PCA ran on the variance stabilized expression data produced as explained above, which together captured the major axes of transcriptomic variation while excluding minor components likely dominated by noise, as done for the multivariate disparity analysis. Plasticity was then quantified for each sibling pair as the multivariate distance between the expression profiles of the two siblings grown under different treatments (dry vs. control). Specifically, for each pair, we calculated the Euclidean distance between their coordinates in the 15-dimensional PCA space in this way:

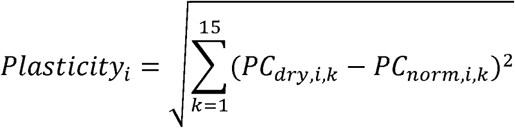

Where *PC_dry,i,k_* and *PC_norm,i,k_* are the scores of the two siblings of pair *i*. on principal component *k*. Differences in transcriptomic plasticity among ploidy levels were then tested using linear mixed-effects models implemented in the lmerTest v.3.1-3 package (Kuznetsova et al. 2017), after evaluating the plasticity values as normally distributed performing a Shapiro–Wilk test. Ploidy was included as a fixed effect and to account for non-independence among related individuals in the same treatment we included family identity (defined by parental cross) as a random intercept.

### Weighted gene co-expression network analysis

We performed weighted gene co-expression network analysis (WGCNA) separately for 30 diploid (2x), 30 neo-tetraploid (neo4x) and 29 established autotetraploid (4x) samples, combining transcriptomes from both treatments in each case, to investigate ploidy-specific patterns of gene co-regulation. The analysis was carried out using the WGCNA v.1.73 package (Langfelder and Horvath 2008) in R v.4.2.2. Firstly, for each ploidy group, we pre-processed the gene expression data creating a DESeqDataSet object using the DESeq2 package (Love et al. 2014), specifying the treatment (drought vs. control) as the design formula. We then estimated size factors using estimateSizeFactors() to normalize for differences in library size. To reduce noise from lowly expressed genes, we retained only those genes with at least 8 raw counts in at least 60% of samples within each dataset. We applied the variance-stabilizing transformation using getVarianceStabilizedData() to obtain homoscedastic expression values suitable for correlation analysis. To further reduce technical noise and improve network structure, we filtered out the 10% least variable genes in each dataset, retaining 16,711 genes in the 2x group, 16,697 in the neo4x group, and 17,175 in the 4x group. The distributions of the variance-stabilized expression values were visually inspected using histograms, boxplots, and quantile–quantile plots to confirm homogeneity across samples, absence of strong outliers and normal distribution.

First, sample clustering using hierarchical clustering detected one outlier sample in the 2x and neo4x datasets there were excluded based on dendrogram inspection, resulting in a consistent number of 29 samples per ploidy group used. A soft-thresholding power (β) of 7 was then selected for each network using the pickSoftThreshold() function. Because raw connectivity measures such as kWithin and kOut are strongly influenced by the choice of soft-thresholding power (β), we used the same β value across all three ploidy-specific networks to ensure comparability. This value was not arbitrarily chosen; rather, it was selected based on standard WGCNA criteria, namely achieving a scale-free topology model fit (R²) ≥ 0.8 while maintaining a reasonable mean connectivity (∼20). This approach ensures that observed differences in network topology reflect biological variation rather than technical artifacts of network construction. We finally used the signed hybrid network type and computed the topological overlap matrix (TOM) with TOMType = “signed”. Gene modules were identified setting a minimum module size of 10 genes and a merge cut height of 0.15.

Finally, to assess the evolutionary persistence of co-expression modules across ploidy levels, we performed module preservation analyses using the function modulePreservation() available in the WGCNA package, with 100 permutations. This approach evaluates whether groups of genes that form modules in the test network retain similar co-expression relationships in another (reference) network, independent of module label assignments. We specifically tested whether and which modules recognised in neo4x (test) are preserved in established tetraploids and diploids networks (references) separately. For each test we selected the strongly preserved modules as the ones showing a Zsummary > 10. The Zsummary statistics integrates measures of module density and connectivity in the test network relative to random expectation and should be interpreted as a measure of retention of co-expression topology.

### Gene Ontology enrichment analysis

To infer the functional involvement of the detected DEGs and the networks’ modules in potential biological processes, we worked with *A. thaliana* orthologs of *A. arenosa* genes obtained using OrthoFinder v.3.0 (Emms and Kelly 2019) and the *A. thaliana* gene universe to perform a Gene Ontology (GO) enrichment test, using the R package topGO v.2.46.0 (Alexa A 2024). We applied the weight01 algorithm with Fisher’s exact test (p < 0.05) as implemented by the package.

## Results

### Phenotypic responses to drought across ploidy levels

To evaluate drought effects at the morphological level for each ploidy, we assessed the effect of drought treatment and ploidy on rosette diameter at the time of RNA collection using linear models (Fig. 1B). Plants exposed to drought developed significantly smaller rosettes compared to controls (Fig. 1B), where neither ploidy (F_2,84_ = 2.09, p = 0.13) nor the treatment × ploidy interaction (F_2,84_ = 1.32, p = 0.27) were statistically significant. We further scored flower number at the end of the experiment as fitness proxy reflecting total investment into reproductive output. We found that it was significantly affected by ploidy (F_2,82_ = 26.74, p = 1.15 × 10^-9^), with established 4x having a considerable higher number of flowers compared to the other two ploidies. Despite an observable trend for decreased flowering in dry conditions (Fig 1B), neither drought treatment (F_2,82_ = 1.04, p = 0.31), nor its interaction with ploidy (F_2,82_ = 0.53, p = 0.59) had a significant effect on the final number of flowers. This indicates that while drought substantially reduced vegetative growth (rosette diameter), being indicative of stress at the time of RNA sampling, it did not significantly impact the plants’ reproductive output (flower number), and that ploidy differences played a stronger role in determining generative traits than in affecting the magnitude of drought sensitivity at the vegetative stage.

**Fig. 1.**
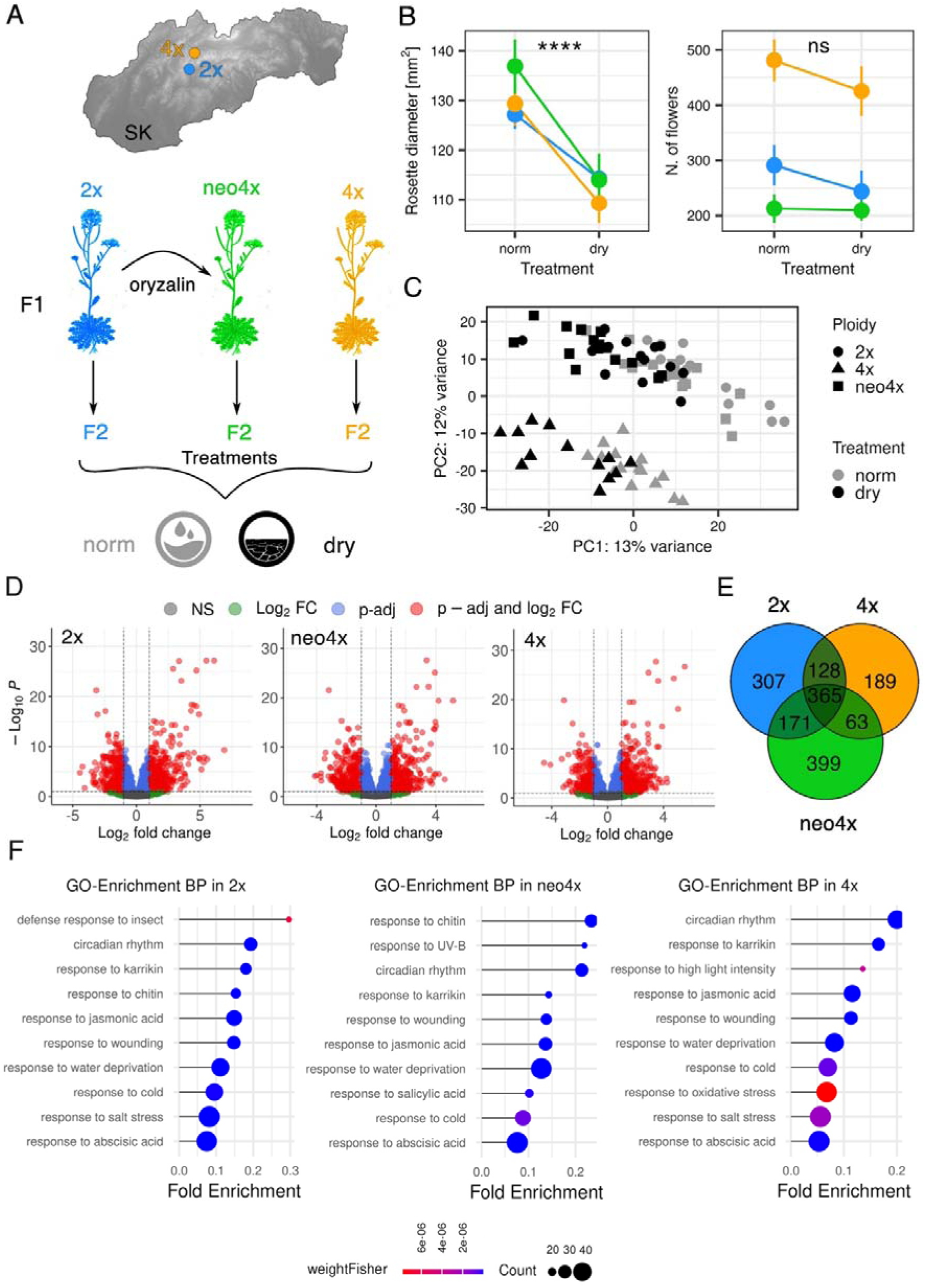
Experimental design, differential expression analysis and gene ontology enrichment. (A) Map of Slovakia showing the location of the two natural *Arabidopsis arenosa* populations used in this study: a diploid population (2x, blue) and a genetically close established autotetraploid population from the same evolutionary lineage and similar habitat (4x, orange). For the experiment, synthetic neo-tetraploids (neo4x, green) were generated by treating diploid F1 seedlings with oryzalin. F2 individuals from all three ploidy types were formed by within-ploidy crossings to reduce maternal effect. They were then grown and subjected to two watering treatments: normal (norm, grey) and water limitation (dry, black). (B) Differences in rosette diameter at vegetative stage (left) and final number of flowers (right) for each ploidy and treatment. Significance of the treatment effect tested by linear model is indicated by asterisks in each plot: rosette diameter F_1,84_ = 24.32, p = 4.06 × 10^-6^; number of flowers F_1,82_ =1.04, p = 0.3107. (C) Principal component analysis (PCA) based on filtered and variance-stabilized gene expression data across all 89 samples, showing a main clustering by treatment (norm vs dry, PC1 explains 13% of variance) and then by ploidy (2x, neo4x, 4x, PC2 explains 12% of variance). (D) Volcano plots showing differentially expressed genes (DEGs) between drought and control conditions within each ploidy group. Genes significantly differentially expressed (adjusted p-value < 0.1 and |log=FC| > 1) are highlighted in red. (E) Venn diagram showing the number of significantly treatment-associated DEGs specific to each ploidy group and their overlap. (F) Ten most significantly enriched Gene Ontology (GO) terms for biological processes (BP) among the significant DEGs in each ploidy group. Bubble plots show the fold enrichment for each enriched GO terms, with dot size proportional to the number of DEGs in the term and colour representing statistical significance (Fisher’s exact test).

### Transcriptomic responses to water limitation vary across ploidy levels

To assess how gene expression and its responses to water limitation differ among diploid (2x), synthetic neo-tetraploid (neo4x), and closely related established autotetraploid (4x) of *A. arenosa*, we analysed fully developed rosette leaf transcriptomes of each ploidy group after subjecting the plants to either control or drought treatments (89 plants in total, 14-15 individuals per ploidy and treatment combination, Fig. 1A). RNA-seq data quality was overall high and consistent across ploidy samples and treatments (Supplementary Table S1). The majority of reads (87.36% on average) mapped uniquely to the reference genome, with low proportions of multi-mapping and unmapped reads. Principal component analysis of filtered and variance-stabilized expression data revealed separation of samples by both treatment and ploidy level, with drought-treated samples generally clustering apart from controls along PC1 and 2x and neo4x samples clustering apart from established 4x along PC2 (Fig. 1C). These patterns are supported by a permutational multivariate analysis of variance (PERMANOVA), where treatment and ploidy interaction explained a significant portion of the variation in global transcriptomic profiles (PERMANOVA: F = 3.76, R² = 0.184, p = 0.001).

Differential expression analysis within each ploidy group identified hundreds of genes responsive to drought (Fig. 1D). The number and identity of differentially expressed genes (DEGs) varied across groups, with neo4x plants showing the highest number of DEGs at the same applied threshold (adjusted p-value < 0.1 and |log=FC| > 1), and 4x plants showing the lowest number of DEGs, especially downregulated, suggesting a distinct transcriptional regulation pattern (Table 1). A substantial core set of 365 genes was differentially expressed in all three ploidies (Fig. 1E), while each group also showed a substantial number of unique DEGs (307 in 2x, 399 in neo4x, and 189 in 4x). Ploidy-specific GO enrichment analysis revealed that drought-responsive genes in all ploidy groups were enriched for relevant biological processes strongly related to drought response, including responses to water deprivation, abscisic acid, salt stress, cold and oxidative stress (Verslues et al. 2006; Osakabe et al. 2014). All groups also shared enrichment for defence-related processes, including responses to jasmonic acid, chitin, and wounding. Indeed, significantly enriched GO-terms inferred for each ploidy type are shared between each ploidy comparison more than what would be expected by chance (Supplementary Fig. S3).

**Table 1.** Number of significantly (adjusted p-value < 0.1 and |log=FC| > 1) differentially expressed genes (DEGs), up and down regulated, for each ploidy type.

| Ploidy | All DEGs | Up DEGs | Down DEGs |
| --- | --- | --- | --- |
| 2x | 971 | 526 | 445 |
| Neo4x | 998 | 485 | 513 |
| 4x | 745 | 483 | 262 |

### Neo-tetraploids show the highest transcriptomic space occupancy

To assess whether neo-tetraploids (neo4x) occupy a broader transcriptomic space than diploids (2x) or established autotetraploids (4x), reflecting a higher occupancy of the phenotypic landscape, we quantified multivariate disparity of each ploidy group as the sum of variances across the first 15 PCs. Disparity differed markedly among groups in both treatments (Fig. 2). The highest dispersion of samples was detected in water deficiency conditions, where neo4x exhibited the highest disparity, followed by 2x and 4x (Fig. 2B, Supplementary Table S2). Pairwise Wilcoxon rank-sum tests revealed significant differences between all ploidies when subjected to drought, while under control treatment disparity was reduced in all ploidies as compared to drought treatment but remained higher in neo4x and 4x (Fig. 2B, Supplementary Table S2). All pairwise differences between ploidies within the normal treatment were significant as well (Supplementary Table S2). Overall, these results indicate that neo4x populations—especially under drought—explore a substantially broader region of transcriptomic space than their diploid progenitors or established autotetraploids, consistent with an expectation of greater variability and/or plasticity in gene expression. To directly test if neo4x higher transcriptomic variability can be framed as plasticity, we calculated the multivariate displacement in PCA space between sibling individuals grown under contrasting water regimes. Therefore, we here define plasticity as the transcriptomic shift in response to environmental change of a genotype. We found a significant effect of ploidy on plasticity magnitude, with neo-tetraploids exhibiting a significantly higher transcriptomic plasticity compared to their ancestral diploids (mean 2x = 121.48, mean neo4x = 147.93, t = 2.19, *p* = 0.039).

**Fig. 2.**
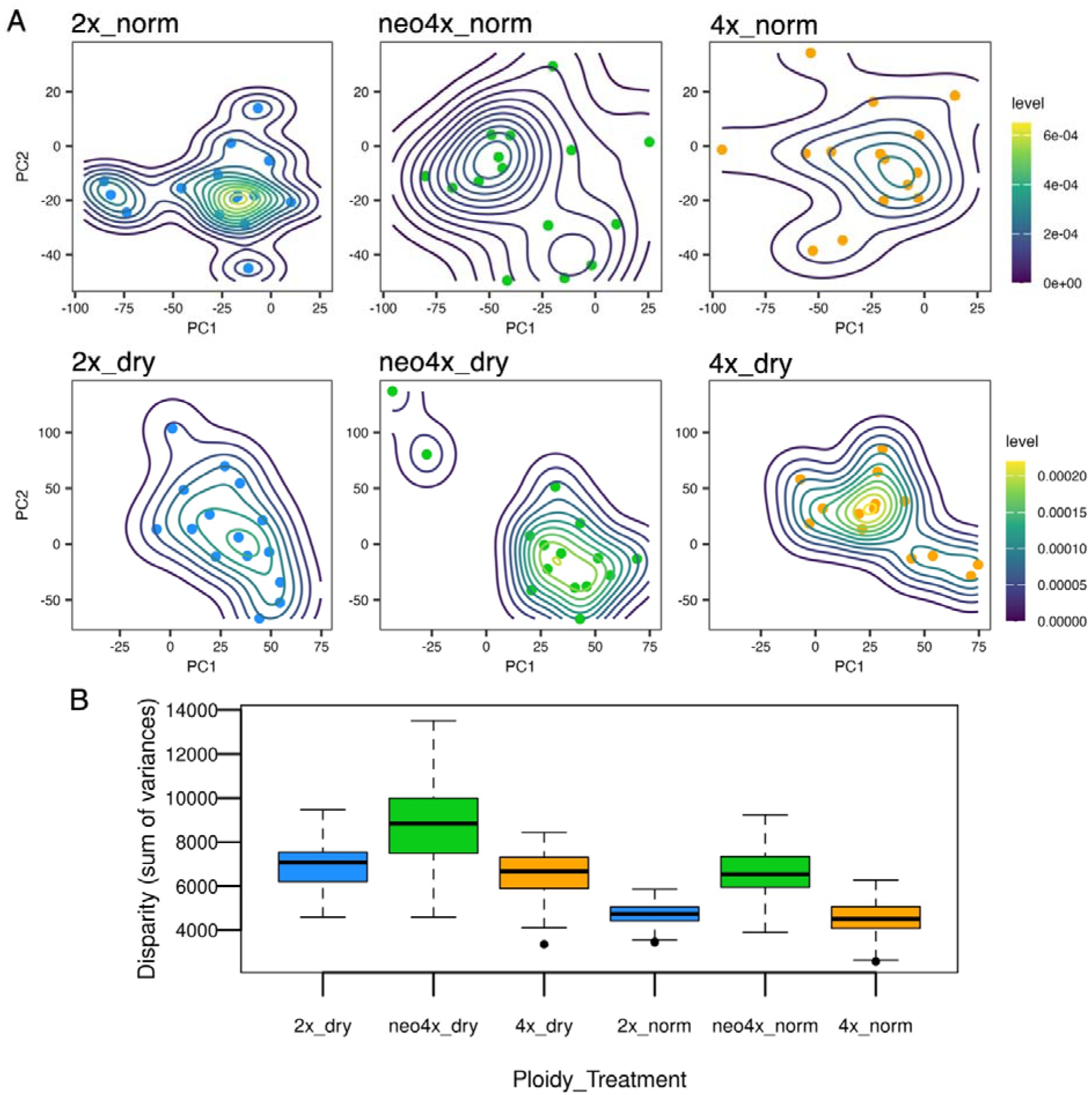
Transcriptomic space occupancy across ploidies and treatments. (A) Principal component analysis (PCA) of variance-stabilized RNA-seq expression data where the separate panels are showing the position of individuals from each ploidy–treatment combination along the first two principal components (explaining 8.4% and 7% of variation, respectively). Density contours represent 2D kernel density estimation of sample distribution in PC1–PC2 space, with individual samples overlaid as points. The higher occupancy of the transcriptomic space by neo4x is here visible as already strongly captured by the first two PCs. (B) Multivariate disparity (sum of variances across the first 15 PCs) calculated for each ploidy–treatment combination using the dispRity package with 1,000 bootstrap replicates. Boxplots show the bootstrap distributions of disparity, indicating the extent of multivariate transcriptomic space occupied by each group.

### Neopolyploid networks show increased fragmentation and inter-modular connectivity

To assess how co-expression network structure differs across ploidy levels, we constructed independent co-expression networks for each ploidy group (including both treatments) using WGCNA. At the same soft-thresholds for all ploidy levels, that ensures comparability, the neo4x network exhibited the highest number of modules (129 before and 115 after merging), nearly double that of the diploid (66/63 modules) and more than the established autotetraploid (86/81 modules) networks (Fig. 3A). Connectivity analyses revealed marked differences in network topology between ploidies. Gene-level intra-modular connectivity (kWithin) was significantly higher in 2x compared to both neo4x and 4x (Fig. 3B). However, this trend loses significance at the modular level, where each ploidy network shows comparable levels of average within-module connectivity (Fig. 3B). Together, these results suggest that while overall module cohesion is similar, diploid networks are characterized by a more uniformly high intra-modular connectivity across genes, whereas neo4x and 4x networks may rely on a smaller number of highly connected nodes, resulting in a less homogeneous internal structure. In contrast, inter-modular connectivity (kOut) was significantly higher in neo4x than in other two ploidy groups at both the gene and module levels (Fig. 3C), indicating that the neo4x network is characterized by greater cross-module interactions. This shift was also reflected in kDiff (kWithin – kOut), where 2x and 4x showed positive values (stronger internal connectivity), while neo4x had lower or near-zero kDiff, particularly at the gene level (Fig. 3D). These patterns indicate that neo-tetraploids exhibit a reorganization of co-expression architecture characterized by reduced intra-modular cohesion and increased inter-modular connectivity. This results in weaker separation between modules and a shift toward a more integrated and less compartmentalized network structure, in contrast to the more clearly modular organization observed in diploids and established autotetraploids (Fig. 3E).

**Fig. 3.**
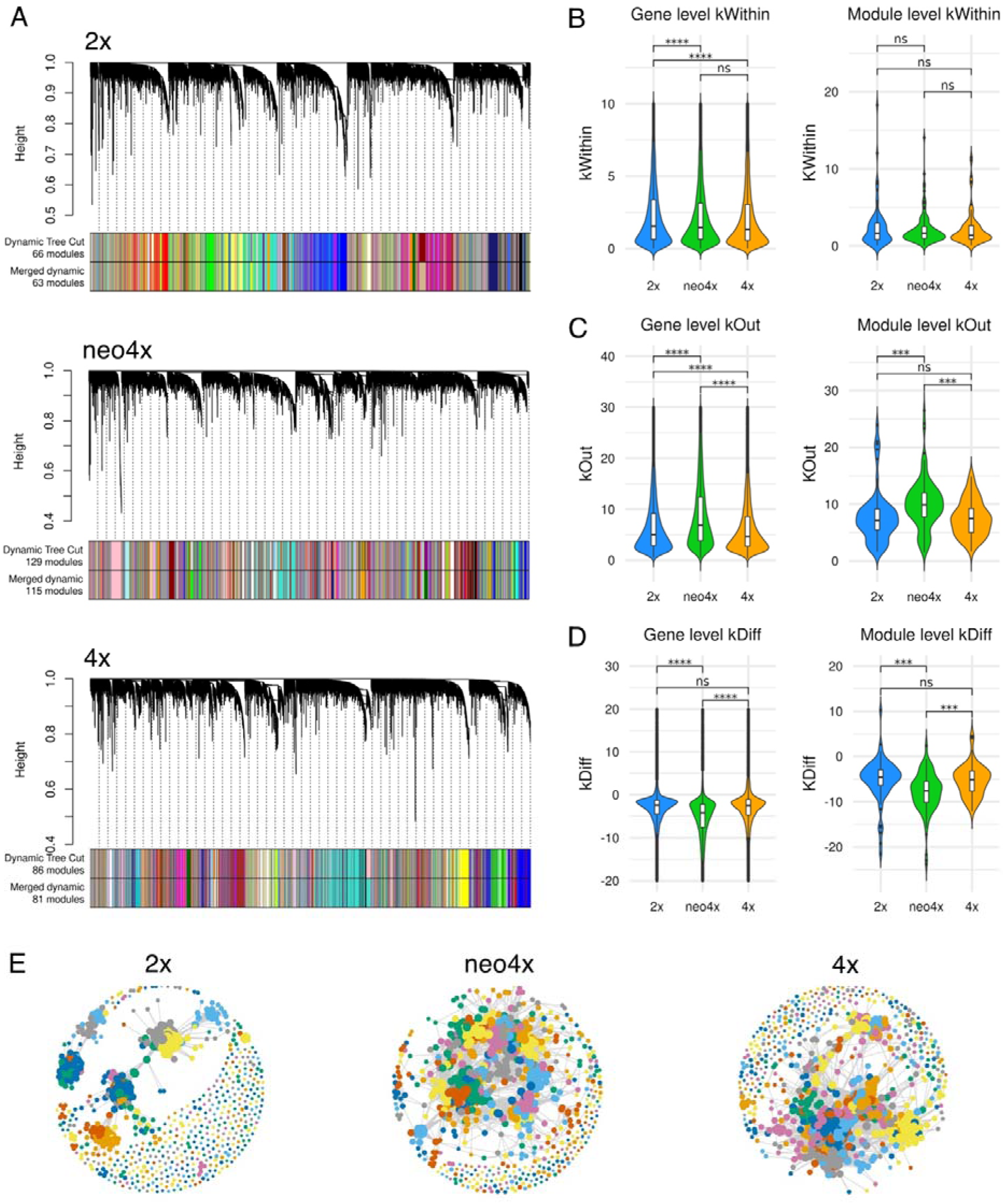
Network modularity and connectivity properties across ploidy levels. (A) Hierarchical clustering dendrograms of gene expression profiles used for co-expression module detection in each ploidy group. Coloured horizontal bars represent modules identified by dynamic tree cut (top row) and after module merging based on eigengene similarity (bottom row). Violin and box plots comparing kWithin (intra-modular connectivity, B), kOut (inter-modular connectivity, C) and kDiff (kWithin – kOut, D) at the gene level (left) and module level (right). Statistical significance was determined using pairwise comparisons on normalised data (****p < 0.0001, ***p < 0.001, ns = not significant). The y-axis limits of each “Gene level” plot were reduced to aid visibility. E) Network visualisation per each ploidy including the 10,000 genes (nodes) with the highest expression variance and the top 5% strongest connections (edges). Nodes are coloured by module assignment, while their size is positively correlated to their connectivity levels.

### Neo-tetraploids show higher plasticity in response to water deficiency stress

We further focused on the ploidy-specific network characteristic in response to water deficiency stress. First, we tested whether transcriptional modules were associated with plants phenotype scored at the time of RNA collection described above. Rosette diameter was indeed significantly correlated with multiple modules in all ploidies, but neo4x displayed the highest number of significant module–trait associations, even after accounting for their larger total number of modules (in 2x, the 11% of modules were significantly correlated to the trait, in neo4x 23% and in 4x 17%; Fig. 4A). This was supported by logistic regression, indicating that neo4x modules were significantly more likely to be correlated with rosette diameter compared to diploid modules (odds ratio ≈ 2.5, *p* = 0.0496), whereas tetraploid modules showed no significant difference from diploids (odds ratio ≈ 1.7, *p* = 0.30). This indicates that the neo4x transcriptome is more extensively linked to plant response to stress, again consistent with an expectation for expanded phenotypic and transcriptomic space in neo-polyploids.

**Figure 4.**
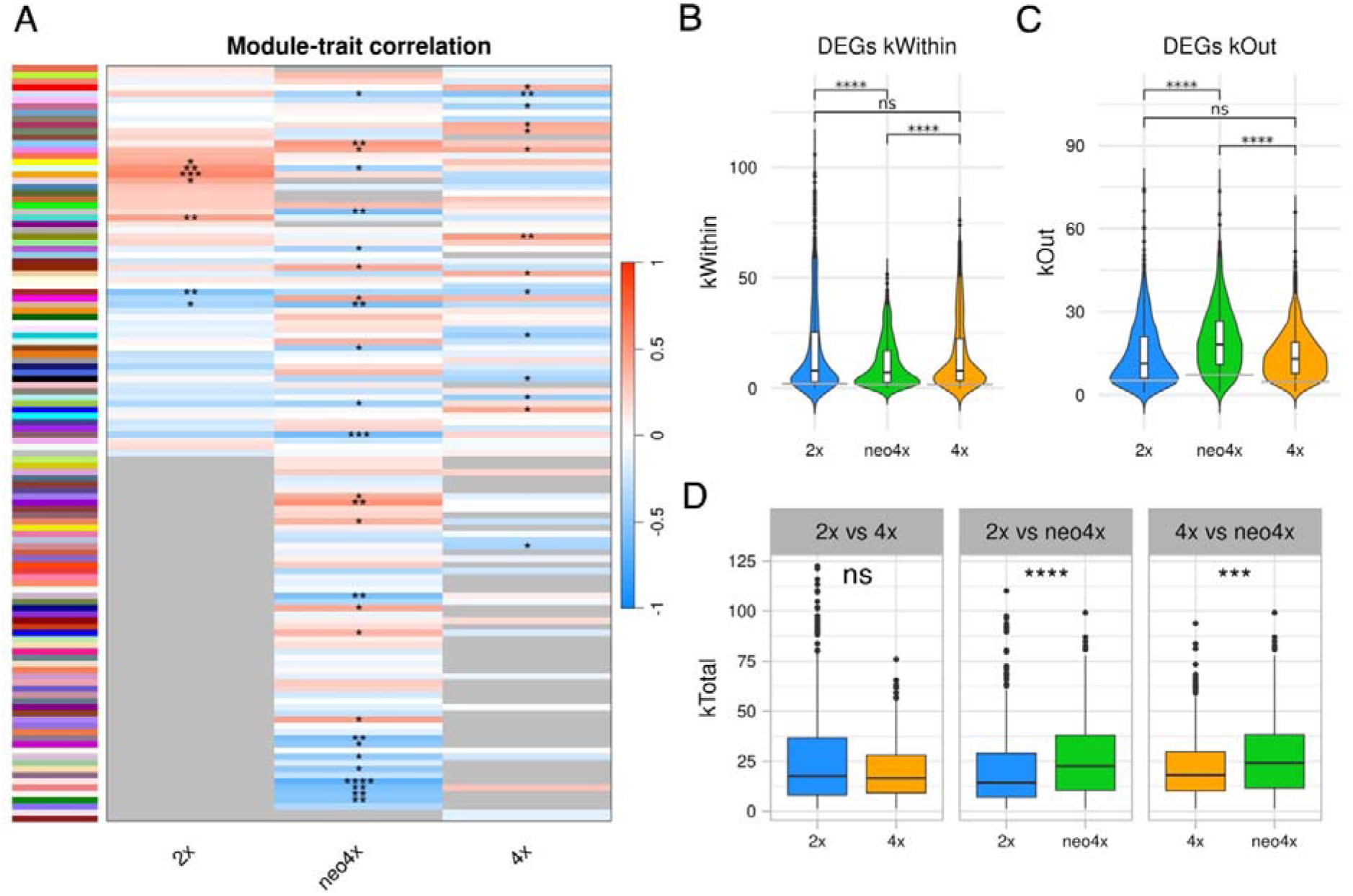
Module–trait correlations and network properties of drought-related differentially expressed genes (DEGs) across ploidy levels. (A) Module–trait correlations between the detected WGCNA modules and rosette diameter. Each row corresponds to a module, reported by its colour name, and blue-red colour scale indicates the strength and direction of the correlation with the phenotype (red = positive, blue = negative). Significant correlations are marked with asterisks (* *p* < 0.05, ** *p* < 0.01, *** *p* < 0.001, **** *p* < 0.0001). Neo4x showed the highest number of modules significantly associated with the trait. Distribution of (B) within-module connectivity (kWithin) and (C) external connectivity (kOut) of DEGs across ploidies. DEGs in neo4x display significantly lower kWithin and significantly higher kOut compared to diploids and tetraploids. The grey segment indicated the median value of the respective kWithin and kOut of the full ploidy-specific network, showcasing a strongly higher connectivity of DEGs compared to the other genes. (D) Total connectivity (kTotal) of ploidy-specific DEGs in each pairwise ploidy comparison. For each pairwise comparison (2x vs 4x, 2x vs neo4x, 4x vs neo4x), only DEGs unique to one ploidy were retained, and their kTotal values compared. While there is no difference in connectivity between 2x-specific and 4x-specific DEGs, neo4x-specific DEGs show significantly higher kTotal than the DEGs specific to 2x and to 4x). Statistical significance in B, C and D was determined using pairwise comparisons on normalised data (\*\*\*\**p* < 0.0001, \*\*\**p* < 0.001, ns = not significant).

We next investigated how differentially expressed genes (DEGs) were integrated within the ploidy-specific co-expression networks. To test whether DEGs are distributed non-randomly across the co-expression modules, we compared the observed counts of DEGs per module to a permutation-based null model that assumes DEGs are distributed proportionally to module size. Across all ploidy levels, the observed distributions did not deviate significantly from the null expectation (2x: χ² = 17490.5, *p* = 0.837; neo4x: χ² = 16074.6 *p* = 0.942; 4x: χ² = 26518.3, *p* = 0.878). These results indicate that in all ploidies the number of modules in which DEGs occur is primarily explained by the relative size of each module, and there is therefore a lack of modules enriched for the drought stress-responsive genes. However, the position of DEGs within the networks differed strongly among ploidies. Across all networks, DEGs were significantly more connected than the average background of non-DEG genes (two-sided permutation test for each ploidy returned *p* = 0), confirming that transcriptional responses tend to affect network hubs rather than peripheral nodes (Fig. 4B–C). Yet, the type of connectivity varied with ploidy. While there was generally no difference in kTotal between ploidies (pairwise comparisons: 2x vs. neo4x, *p.adj* = 0.462; 2x vs. 4x, *p.adj* = 1.000; neo4x vs. 4x, *p.adj* = 0.597), in neo4x DEGs showed markedly higher kOut (external connectivity to other modules) but lower kWithin (intra-module connectivity) compared to diploids and tetraploids (Fig. 4B–C). While this pattern reflects the general trend described for the full network, it suggests that the hub nodes in neo4x coordinate the response to stress as “inter-modular connectors” or network bridges rather than being embedded within tightly co-expressed modules, like in the other two ploidies. Our results about transcriptomic plasticity, together with such topology suggest that in neo-polyploids stress-responsive genes are potentially facilitating broader regulatory flexibility and phenotypic plasticity.

To investigate a ploidy-specific response to stress, we examined ploidy-specific DEGs (differentially expressed genes in one ploidy, non-overlapping between pairs of ploidies) and found that those unique to neo-polyploids were more central (higher kTotal) than DEGs unique to diploids or established tetraploids, further underscoring that neo-polyploids rely on new pathways that tend to affect a higher number of phenotypic traits (Fig. 4D). No difference was observed between diploid-and tetraploid-specific DEGs, suggesting that tetraploid would evolve to reach a diploid-like state of connectivity in the long-term (Fig. 4D).

### A subset of novel neo-tetraploid modules is retained in natural tetraploids

Finally, we explored whether the novel regulatory structures we detected as triggered by WGD *per se* were retained during following tetraploid’s evolution. Performing a module preservation analysis (*Z* summary > 10), we identified 70 neo4x modules preserved from the 2x network, and 77 neo4x modules preserved in the 4x network (Fig. 5). Within these, 12 neo4x modules showed significant preservation in the 4x network, while not being preserved in the diploid one (Fig. 5). This indicates that novel co-expression structures originating from WGD have the potential to prove adaptive and stabilise to be retained long-term. Our result indeed showed that genes belonging to the 12 preserved neo4x modules (altogether 713 genes) were often distributed across multiple 4x modules (74 modules in total, with each preserved neo4x module distributed in 21 4x modules on average). Nevertheless, preservation statistics indicated that these genes maintained non-random co-expression connectivity patterns in the 4x network (mean *Z* summary = 12.77), demonstrating overall retention of the WGD-triggered network topology, but with further adjustment into distinct modules in established 4x (Fig. 5). Moreover, genes belonging to the 12 preserved modules were significantly enriched for biological processes important for drought stress tolerance and water retention (also found for example in Zhou et al. 2019; Wang, Fan, et al. 2022 and Xu et al. 2023), such as cell wall and extracellular structure organization, including cuticle and cutin-based cuticle development and polysaccharide and β-glucan biosynthesis (Supplementary Table S3). Interestingly, constitutive between-ploidy differences in drought-associated genes have been observed also in previous differential expression analyses in other *A. arenosa* populations (Srikant et al. 2024). Although this is an interesting association, we acknowledge that a direct test of fitness effect of such post-WGD preserved modules would be needed to prove adaptive value of such changes.

**Figure 5.**
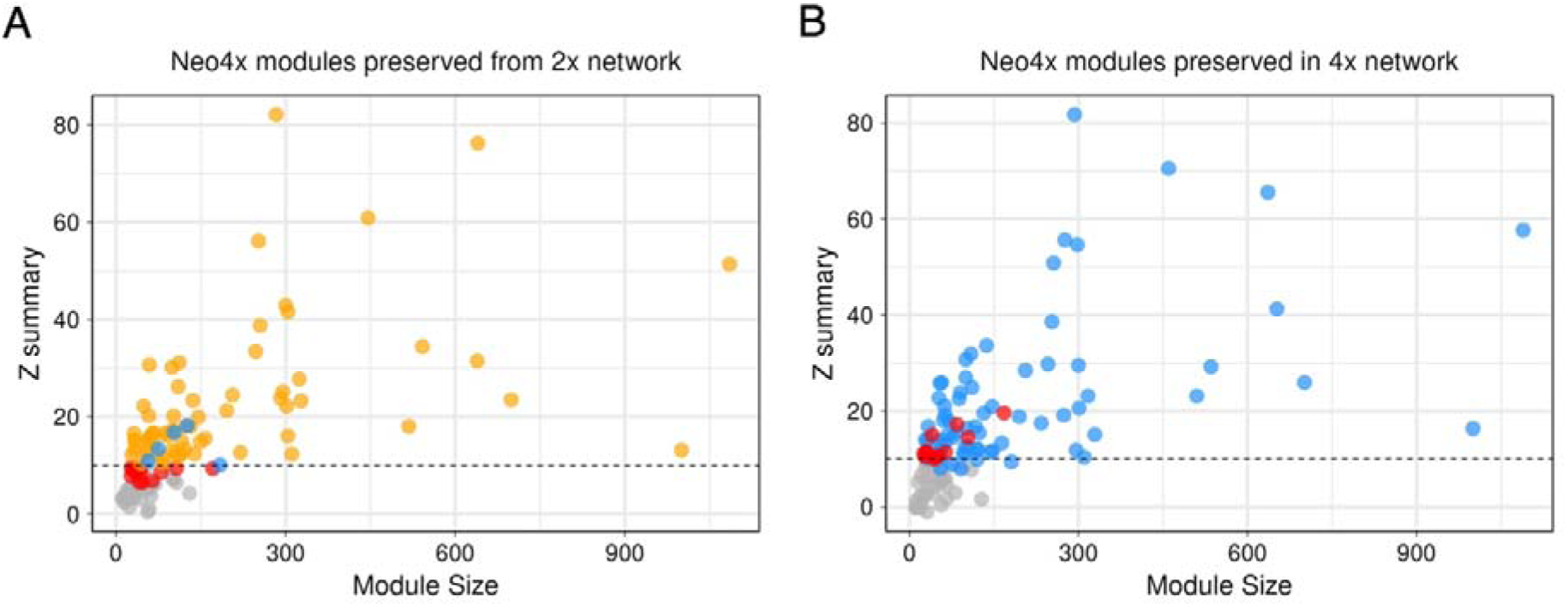
Differential preservation of neo-tetraploid co-expression modules in diploid and established tetraploid networks. (A) Preservation of neo4x co-expression modules in the diploid (2x) network and (B) preservation of neo4x modules in the established tetraploid (4x) network, assessed using the WGCNA module preservation statistic (*Z* summary). Each point represents a neo4x module, plotted according to its size (number of genes) and preservation score. The dashed horizontal line indicates the threshold for strong preservation (*Z* summary = 10), with modules above this value considered highly preserved. Grey points represent modules that did not reach the preservation threshold in the respective comparison. In panel A, orange points indicate neo4x modules that are also strongly preserved in the 4x network, while blue points indicate modules that are highly preserved only in the 2x networks. In panel B, blue points indicate modules that are also highly preserved in 2x network. In both panels red points represent the 12 modules that are strongly preserved only in the established tetraploid network and not in the diploid one.

## Discussion

Polyploids have long been viewed through the lens of Goldschmidt’s “hopeful monsters” theoretical concept (Otto and Whitton 2000), where sudden saltational genomic reconfigurations have the potential to generate novel and advantageous phenotypes, but also carry the risk of maladaptive instability (Goldschmidt 1941). However, while polyploidy is pervasive in nature and especially in plant evolution (Otto and Whitton 2000; Crow and Wagner 2006; Wendel 2015; Van de Peer et al. 2017), its short and long-term consequences for fitness, gene regulation, and adaptability remain debated (Comai 2005; Spoelhof et al. 2017; Fox et al. 2020; Clo 2022). Are neo-polyploids genuinely “hopeful monsters,” i.e. having transiently expanded phenotypic possibilities compared to their ancestor, while still maintaining a high fitness (Parisod 2024)?

Our results provide an empirical step forward in bridging this gap. By comparing diploids, synthetic neo-tetraploids, and established autotetraploids of *Arabidopsis arenosa*, we show that the neo-tetraploid state is indeed characterized by a transient phase of transcriptomic expansion, higher plasticity and network rewiring. In line with previous theoretical computational work (Yao et al. 2019; Ebadi et al. 2023), we show that polyploidization *per se* enhances network complexity and phenotypic (transcriptional) variance. We indeed observe that neo-tetraploid plants have a higher number of differentially expressed genes under stress, occupy a broader transcriptomic space and display a co-expression architecture dominated by a higher number of smaller modules tightly inter-connected, suggesting enhanced regulatory flexibility. Importantly, this increased connectivity is directly associated with plant performance, as evidenced by the larger fraction of modules correlated with a vegetative fitness proxy (rosette diameter) in neo-tetraploids. Over evolutionary time, however, overall patterns of gene expression appear to undergo compensatory changes and return to a diploid-like state—i.e. following an evolutionary trajectory of Type A characters (Bomblies 2020): established autotetraploids recover relatively more modular, diploid-like structures, accompanied by reduced transcriptomic disparity. Similar evolutionary trajectory was recently described for constitutive differential expression in other *A. arenosa* populations (Srikant et al. 2024). Our results thus document that such evolutionary trends are likely general pattern in *A. arenosa*.

From an underlying mechanistic point of view, whole-genome duplication is widely recognized to induce a transcriptional shock (Comai 2005; Doyle and Coate 2019), disrupting pre-existing regulatory balances and resulting in altered gene expression and co-expression patterns. In our study, this is most evident in neo-tetraploids, where gene networks are more fragmented and stress-responsive genes tend to act as inter-modular connectors rather than within modules hubs. Modularity, i.e. the occurrence of many modules of tightly co-expressed genes relatively poorly connected to external nodes, is expected to facilitate evolutionary adjustments without perturbing the balance of the full system (Davidson 2010; Espinosa-Soto 2018). In other words, it confers robustness while also enabling evolvability (Kashtan and Alon 2005; Van Belleghem et al. 2017). We observe that after polyploidization the transcriptional shock leads to a reduction in modularity, therefore possibly negatively affecting robustness and evolvability. However, we also found an increased number of modules, similarly to what has been observed for GRNs motifs after ancient WGDs in angiosperms (Almeida-Silva and Van de Peer 2023). We hypothesize that these characteristics could simultaneously expand the range of possible regulatory interactions, potentially exposing novel transcriptional and phenotypic configurations that allow to further explore the fitness landscape (Yao et al. 2019; Ebadi et al. 2023; Parisod 2024). Enlarged phenotypic variation is manifested in our data as larger number of differentially expressed genes and broader transcriptomic landscape in multivariate analyses of neo-tetraploids, especially under stress. This window of regulatory flexibility increases the likelihood that at least some of the newly expressed phenotypes will prove advantageous under stress and changing environments.

In the long term such network topology is likely to be maladaptive (Conant and Wolfe 2006; Whitacre and Bender 2010; Srikant et al. 2024), although we have not observed any immediate fitness cost of WGD *per se* neither in terms of growth nor flower production. In line with this, established autotetraploids of *A. arenosa* seem to have re-stabilized their transcriptomic architecture, reverting towards a diploid-like level of modularity. However, they might have retained a subset of the variation that arose during the neo-polyploid shock that turned out to be beneficial for fitness. Indeed, in our experiment established tetraploids preserved WGD-specific co-expression structures encompassing genes with functions relevant in the context of drought stress. Retention of favourable variants arising from WGD in longer term (at the level of diploidizing genomes) is indeed commonly observed (Otto and Whitton 2000; Wu et al. 2020; Almeida-Silva and Van de Peer 2023), e.g. for cold-related genes in *Brassicaceae* (Song et al. 2020), genes involved in signal transduction and transcription in *Arabidopsis thaliana* (Blanc and Wolfe 2004), genes involved in adaptation to salinity in mangroves (Xu et al. 2023), and salt and heavy metal responsive genes in *Cakile maritima* (Thomas et al. 2024). Our results complement this growing body of evidence by suggesting that a some of the initial post-WGD transcriptomic variation can be fixed early, already during polyploid establishment phase.

Despite topological differences between co-expression networks of different ploidies, stress-responsive differentially expressed genes (DEGs) were consistently more central than non-DEGs across all ploidies, indicating that transcriptional responses to stress preferentially target hub genes. This suggests that drought adaptation in the long-term might rely on reconfiguring key regulatory nodes rather than peripheral genes, a finding in line with previous studies across various diploid taxa and different conditions (Wang et al. 2010; Frachon et al. 2017; Hämälä et al. 2020; Rennison and Peichel 2022; Celestini et al. 2025). However, ploidy-specific DEGs highlighted important differences: neo4x-specific DEGs were significantly more central than those unique to diploids or established tetraploids, meaning they likely affect a higher number of traits. While the specific mechanisms leading to this pattern are unclear, we hypothesize that, similarly to what explained above, this might allow for jumping in the fitness landscape and quickly landing on a fitness peak.

Finally, several limitations of our study should be acknowledged. First, our comparisons were based on a single natural population per ploidy, which limits generalisation of our findings. However, our data are based on large and genetically variable population sample (∼12 families per ploidy including the neo-polyploid cohort). In addition, overall patterns of ploidy-related transcriptomic differences found in our sample are in line with findings from other *A. arenosa* populations and tissues (Srikant et al. 2024). Second, although we cannot completely exclude that the effect of local differences in drought adaptation between natural populations can contribute to the observed ploidy-related differences, our sampling was focused to minimize such a risk. Given the close geographic position, genetic relatedness and similar niche (Monnahan et al. 2019; Wos et al. 2019; Morgan et al. 2020) of our diploid and tetraploid populations, we consider unlikely that our results are driven by pre-existing ecological divergence rather than ploidy level *per se*. Third, our transcriptomic analyses were restricted to a single time point and one tissue type, which necessarily provides only a partial view of the drought response and may overlook temporal or organ-specific regulatory dynamics. However, such limited scope shall rather lead to underestimation of the differences observed here. Finally, we acknowledge that our experiment did not reveal a clear performance advantage of any ploidy under the applied drought regime (at least in terms of reproductive allocation or vegetative output), and therefore our results should not be interpreted as direct evidence of polyploid superiority under stress. Rather, our findings demonstrate that WGD and subsequent evolution reshape transcriptional regulation and network architecture, thereby providing alternative routes to stress response. Future experiments that extend to multiple populations and more detailed mechanistic insights into transcriptomic responses to stress will be necessary to clarify under which ecological contexts the transcriptional instability of neo-polyploids promotes novel adaptive opportunities.

## Conclusions

Our results demonstrate that whole-genome duplication *per se* leads to significant perturbation in gene expression patterns, particularly in the architecture of co-expression networks. Although the overall network architecture later stabilizes in established tetraploids, some of the WGD-associated changes may be retained, possibly providing adaptive benefit in particular environmental contexts. Overall, our study supports the notion that neo-polyploids embody a transient “hopeful monster” state — a window of transcriptomic expansion and increased cross-module connectivity, consistent with increased regulatory flexibility and expanded phenotypic potential.

## Acknowledgements

We thank Veronika Vlčková for assistance with plant cultivations and Jana Nosková for laboratory work. This work was supported by the Czech Science Foundation (project No 26-23167S led by F.K) and the European Research Council (ERC) under the European Union’s Horizon 2020 research and innovation programme (ERC-StG 850852 DOUBLE ADAPT to FK). Additional support was provided by the Czech Academy of Sciences (long-term research development project No. RVO 67985939). Computational resources were provided by the e-INFRA CZ project (ID:90254), supported by the Ministry of Education, Youth and Sports of the Czech Republic.

## References

Alexa A RJ. 2024. topGO: Enrichment Analysis for Gene Ontology. Available from: http://bioconductor.org/packages/topGO/

Alix K, Gérard PR, Schwarzacher T, Heslop-Harrison JSP. 2017. Polyploidy and interspecific hybridization: partners for adaptation, speciation and evolution in plants. Ann. Bot. 120:183–194.

Allario T, Brumos J, Colmenero-Flores JM, Tadeo F, Froelicher Y, Talon M, Navarro L, Ollitrault P, Morillon R. 2011. Large changes in anatomy and physiology between diploid Rangpur lime (Citrus limonia) and its autotetraploid are not associated with large changes in leaf gene expression. J. Exp. Bot. 62:2507–2519.

Almeida-Silva F, Van de Peer Y. 2023. Whole-genome duplications and the long-term evolution of gene regulatory networks in angiosperms. Mol. Biol. Evol. 40:msad141.

Andrews S. 2010. FastQC: a quality control tool for high throughput sequence data.

Aversano R, Scarano M-T, Aronne G, Caruso I, D’Amelia V, De Micco V, Fasano C, Termolino P, Carputo D. 2015. Genotype-specific changes associated to early synthesis of autotetraploids in wild potato species. Euphytica 202:307–316.

Bates D, Mächler M, Bolker B, Walker S. 2015. Fitting Linear Mixed-Effects Models Using lme4. J. Stat. Softw. 67:1–48.

Blanc G, Wolfe KH. 2004. Functional divergence of duplicated genes formed by polyploidy during Arabidopsis evolution. Plant Cell 16:1679–1691.

Blighe K, Rana S, Lewis M. 2022. EnhancedVolcano: Publication-ready volcano plots with enhanced colouring and labeling. Available from: https://github.com/kevinblighe/EnhancedVolcano

Bohutínská M, Petříková E, Booker TR, Vives Cobo C, Vlček J, Šrámková G, Poupětová A, Hojka J, Marhold K, Yant L, et al. 2024. Polyploids broadly generate novel haplotypes from trans-specific variation in Arabidopsis arenosa and Arabidopsis lyrata. PLoS Genet. 20:e1011521.

Bomblies K. 2020. When everything changes at once: finding a new normal after genome duplication. Proc. Biol. Sci. 287:20202154.

Bouzid M, He F, Schmitz G, Häusler RE, Weber APM, Mettler-Altmann T, De Meaux J. 2019. Arabidopsis species deploy distinct strategies to cope with drought stress. Ann. Bot. 124:27–40.

Bramsiepe J, Krabberød AK, Bjerkan KN, Alling RM, Johannessen IM, Hornslien KS, Miller JR, Brysting AK, Grini PE. 2023. Structural evidence for MADS-box type I family expansion seen in new assemblies of Arabidopsis arenosa and A. lyrata. Plant J. 116:942–961.

Celestini S, Duchoslav M, Nezamivand-Chegini M, Gerchen J, Šrámková G, Wijfjes R, Krejčová A, Kuzmanović N, Španiel S, Schneeberger K, et al. 2025. Genomic basis of adaptation to serpentine soil in two Alyssum species shows convergence with Arabidopsis across 20 million years of divergence. Ann. Bot.:mcaf141.

Chao D-Y, Dilkes B, Luo H, Douglas A, Yakubova E, Lahner B, Salt DE. 2013. Polyploids exhibit higher potassium uptake and salinity tolerance in Arabidopsis. Science 341:658–659.

Church SA, Spaulding EJ. 2009. Gene expression in a wild autopolyploid sunflower series. J. Hered. 100:491–495.

Clo J. 2022. Polyploidization: Consequences of genome doubling on the evolutionary potential of populations. Am. J. Bot. 109:1213–1220.

Comai L. 2005. The advantages and disadvantages of being polyploid. Nat. Rev. Genet. 6:836–846.

Conant GC, Wolfe KH. 2006. Functional partitioning of yeast co-expression networks after genome duplication. PLoS Biol. 4:e109.

Crow KD, Wagner GP. 2006. What Is the Role of Genome Duplication in the Evolution of Complexity and Diversity? Mol. Biol. Evol. 23:887–892.

Cuypers TD, Hogeweg P. 2014. A synergism between adaptive effects and evolvability drives whole genome duplication to fixation. PLoS Comput. Biol. 10:e1003547.

Davidson EH. 2010. The regulatory genome: gene regulatory networks in development and evolution. Elsevier

De Smet R, Van de Peer Y. 2012. Redundancy and rewiring of genetic networks following genome-wide duplication events. Curr. Opin. Plant Biol. 15:168–176.

Dobin A, Gingeras TR. 2015. Mapping RNA-seq reads with STAR. Curr. Protoc. Bioinformatics 51:11.14.1–11.14.19.

Doležel J, Greilhuber J, Suda J. 2007. Estimation of nuclear DNA content in plants using flow cytometry. Nat. Protoc. 2:2233–2244.

Doyle JJ, Coate JE. 2019. Polyploidy, the nucleotype, and novelty: The impact of genome doubling on the biology of the cell. Int. J. Plant Sci. 180:1–52.

Duan T, Sicard A, Glémin S, Lascoux M. 2023. Expression pattern of resynthesized allotetraploid Capsella is determined by hybridization, not whole-genome duplication. New Phytol. 237:339–353.

Ebadi M, Bafort Q, Mizrachi E, Audenaert P, Simoens P, Van Montagu M, Bonte D, Van de Peer Y. 2023. The duplication of genomes and genetic networks and its potential for evolutionary adaptation and survival during environmental turmoil. Proc. Natl. Acad. Sci. 120:e2307289120.

Emms DM, Kelly S. 2019. OrthoFinder: phylogenetic orthology inference for comparative genomics. Genome Biol. 20:238.

Espinosa-Soto C. 2018. On the role of sparseness in the evolution of modularity in gene regulatory networks. PLoS Comput. Biol. 14:e1006172.

Ewels PA, Magnusson M, Lundin S, Käller M. 2016. MultiQC: summarize analysis results for multiple tools and samples in a single report. Bioinformatics 32:3047–3048.

Fox DT, Soltis DE, Soltis PS, Ashman T-L, Van de Peer Y. 2020. Polyploidy: A Biological Force From Cells to Ecosystems. Trends Cell Biol. 30:688–694.

Frachon L, Libourel C, Villoutreix R, Carrère S, Glorieux C, Huard-Chauveau C, Navascués M, Gay L, Vitalis R, Baron E, et al. 2017. Intermediate degrees of synergistic pleiotropy drive adaptive evolution in ecological time. *Nat*. Ecol. Evol. 1:1551–1561.

Fusco D, Grassi L, Bassetti B, Caselle M, Cosentino Lagomarsino M. 2010. Ordered structure of the transcription network inherited from the yeast whole-genome duplication. BMC Syst. Biol. 4:77.

Gao R, Wang H, Dong B, Yang X, Chen S, Jiang J, Zhang Z, Liu C, Zhao N, Chen F. 2016. Morphological, genome and gene expression changes in newly induced autopolyploid Chrysanthemum lavandulifolium (Fisch. Ex trautv.) makino. Int. J. Mol. Sci. 17:1690.

Goldschmidt R. 1941. The material basis of evolution. Philosophy of Science [Internet]. Available from: 10.2307/1437764

Guggisberg A, Liu X, Suter L, Mansion G, Fischer MC, Fior S, Roumet M, Kretzschmar R, Koch MA, Widmer A. 2018. The genomic basis of adaptation to calcareous and siliceous soils in Arabidopsis lyrata. Mol. Ecol. 27:5088–5103.

Guillerme T. 2018. dispRity: A modular R package for measuring disparity. Methods in Ecology and Evolution 9:1755–1763.

Hämälä T, Gorton AJ, Moeller DA, Tiffin P. 2020. Pleiotropy facilitates local adaptation to distant optima in common ragweed (Ambrosia artemisiifolia). PLoS Genet. 16:e1008707.

Kashtan N, Alon U. 2005. Spontaneous evolution of modularity and network motifs. Proc. Natl. Acad. Sci. U. S. A. 102:13773–13778.

Knotek A, Konečná V, Wos G, Požárová D, Šrámková G, Bohutínská M, Zeisek V, Marhold K, Kolář F. 2020. Parallel alpine differentiation in Arabidopsis arenosa. Front. Plant Sci. 11:561526.

Kolář F, Čertner M, Suda J, Schönswetter P, Husband BC. 2017. Mixed-ploidy species: progress and opportunities in polyploid research. 22:1041–1055.

Kolář F, Lučanová M, Záveská E, Fuxová G, Mandáková T, Španiel S, Senko D, Svitok M, Kolník M, Gudžinskas Z, et al. 2016. Ecological segregation does not drive the intricate parapatric distribution of diploid and tetraploid cytotypes of the Arabidopsis arenosa group (Brassicaceae). Biol. J. Linn. Soc. Lond. 119:673–688.

Konečná V, Bray S, Vlček J, Bohutínská M, Požárová D, Choudhury RR, Bollmann-Giolai A, Flis P, Salt DE, Parisod C, et al. 2021. Parallel adaptation in autopolyploid Arabidopsis arenosa is dominated by repeated recruitment of shared alleles. Nat. Commun. 12:4979.

Kuznetsova A, Brockhoff PB, Christensen RHB. 2017. LmerTest package: Tests in linear mixed effects models. J. Stat. Softw. 82:1–26.

Langfelder P, Horvath S. 2008. WGCNA: an R package for weighted correlation network analysis. BMC Bioinformatics 9:559.

Liu S, Yang Y, Wei F, Duan J, Braynen J, Tian B, Cao G, Shi G, Yuan J. 2017. Autopolyploidy leads to rapid genomic changes in Arabidopsis thaliana. Theory Biosci. 136:199–206.

Love MI, Huber W, Anders S. 2014. Moderated estimation of fold change and dispersion for RNA-seq data with DESeq2. Genome Biol. 15:550.

Monnahan P, Kolář F, Baduel P, Sailer C, Koch J, Horvath R, Laenen B, Schmickl R, Paajanen P, Šrámková G, et al. 2019. Pervasive population genomic consequences of genome duplication in Arabidopsis arenosa. Nat Ecol Evol 3:457–468.

Morgan EJ, Čertner M, Lučanová M, Kubíková K, Marhold K, Kolář F. 2020. Niche similarity in diploid=autotetraploid contact zones of Arabidopsis arenosa across spatial scales. Am. J. Bot. 107:1375–1388.

Oksanen J, Blanchet FG, Kindt R, Legendre P, Minchin PR, O’hara RB. 2025. Vegan: Community ecology package. Available from: https://vegandevs.github.io/vegan/

Osakabe Y, Osakabe K, Shinozaki K, Tran L-SP. 2014. Response of plants to water stress. Front. Plant Sci. 5:86.

Otto SP, Whitton J. 2000. Polyploid incidence and evolution. Annu. Rev. Genet. 34:401–437.

Padilla-García N, Šrámková G, Záveská E, Šlenker M, Clo J, Zeisek V, Lučanová M, Rurane I, Kolář F, Marhold K. 2023. The importance of considering the evolutionary history of polyploids when assessing climatic niche evolution. J. Biogeogr. 50:86–100.

Parisod C. 2024. Duplicated gene networks promote “hopeful” phenotypic variation. Trends Genet. 40:109–111.

Parisod C, Holderegger R, Brochmann C. 2010. Evolutionary consequences of autopolyploidy: Research review. New Phytol. 186:5–17.

Peterson RA, Cavanaugh JE. 2020. Ordered quantile normalization: a semiparametric transformation built for the cross-validation era. J. Appl. Stat. 47:2312–2327.

Preite V, Sailer C, Syllwasschy L, Bray S, Ahmadi H, Krämer U, Yant L. 2019. Convergent evolution in Arabidopsis halleri and Arabidopsis arenosa on calamine metalliferous soils. Philos. Trans. R. Soc. Lond. B Biol. Sci. 374:20180243.

Rennison DJ, Peichel CL. 2022. Pleiotropy facilitates parallel adaptation in sticklebacks. Mol. Ecol. 31:1476–1486.

Rice A, Glick L, Abadi S, Einhorn M, Kopelman NM, Salman-Minkov A, Mayzel J, Chay O, Mayrose I. 2015. The Chromosome Counts Database (CCDB) - a community resource of plant chromosome numbers. New Phytol. 206:19–26.

Rosellini D, Ferradini N, Allegrucci S, Capomaccio S, Zago ED, Leonetti P, Balech B, Aversano R, Carputo D, Reale L, et al. 2016. Sexual polyploidization in Medicago sativa L.: Impact on the phenotype, gene transcription, and genome methylation. G3 (Bethesda) 6:925–938.

Santoro DF, Anderson AW, Alavi SN, Malatesta Pierleoni VA, Rosellini D. 2025. Whole genome duplication drives transcriptome reprogramming in response to drought in alfalfa. Plant Cell Rep. 44:209.

Santoro DF, Marconi G, Capomaccio S, Bocchini M, Anderson AW, Finotti A, Confalonieri M, Albertini E, Rosellini D. 2025. Polyploidization-driven transcriptomic dynamics in Medicago sativa neotetraploids: mRNA, smRNA and allele-specific gene expression. BMC Plant Biol. 25:108.

Scarrow M, Wang Y, Sun G. 2021. Molecular regulatory mechanisms underlying the adaptability of polyploid plants. Biol. Rev. Camb. Philos. Soc. 96:394–407.

Scott AL, Richmond PA, Dowell RD, Selmecki AM. 2017. The influence of polyploidy on the evolution of yeast grown in a sub-optimal carbon source. Mol. Biol. Evol. 34:2690–2703.

Selmecki AM, Maruvka YE, Richmond PA, Guillet M, Shoresh N, Sorenson AL, De S, Kishony R, Michor F, Dowell R, et al. 2015. Polyploidy can drive rapid adaptation in yeast. Nature 519:349–352.

Soltis PS, Soltis DE. 2000. The role of genetic and genomic attributes in the success of polyploids. Proc. Natl. Acad. Sci. U. S. A. 97:7051–7057.

Song X-M, Wang J-P, Sun P-C, Ma X, Yang Q-H, Hu J-J, Sun S-R, Li Y-X, Yu J-G, Feng S-Y, et al. 2020. Preferential gene retention increases the robustness of cold regulation in Brassicaceae and other plants after polyploidization. Hortic. Res. 7:20.

Spoelhof JP, Soltis PS, Soltis DE. 2017. Pure polyploidy: Closing the gaps in autopolyploid research: Pure polyploidy. J. Syst. Evol. 55:340–352.

Srikant T, Gonzalo A, Bomblies K. 2024. Chromatin accessibility and gene expression vary between a new and evolved autopolyploid of Arabidopsis arenosa. Mol. Biol. Evol. 41:msae213.

Stupar R, Bhaskar P, Yandell B, Rensink W, Hart A, Shu O, Veilleux R, Busse JS, Erhardt RJ, Buell CR, et al. 2007. Phenotypic and Transcriptomic Changes Associated With Potato Autopolyploidization. Genetics 176:2055–2067.

Temsch EM, Koutecký P, Urfus T, Šmarda P, Doležel J. 2022. Reference standards for flow cytometric estimation of absolute nuclear DNA content in plants. Cytometry A 101:710–724.

Thomas SK, Hoek KV, Ogoti T, Duong H, Angelovici R, Pires JC, Mendoza-Cozatl D, Washburn J, Schenck CA. 2024. Halophytes and heavy metals: A multi-omics approach to understand the role of gene and genome duplication in the abiotic stress tolerance of Cakile maritima. Am. J. Bot. 111:e16310.

Tossi VE, Martínez Tosar LJ, Laino LE, Iannicelli J, Regalado JJ, Escandón AS, Baroli I, Causin HF, Pitta-Álvarez SI. 2022. Impact of polyploidy on plant tolerance to abiotic and biotic stresses. Front. Plant Sci. 13:869423.

Van Belleghem SM, Rastas P, Papanicolaou A, Martin SH, Arias CF, Supple MA, Hanly JJ, Mallet J, Lewis JJ, Hines HM, et al. 2017. Complex modular architecture around a simple toolkit of wing pattern genes. Nat. Ecol. Evol. 1:52.

Van de Peer Y, Ashman T, Soltis P, Soltis D. 2020. Polyploidy: an evolutionary and ecological force in stressful times. Plant Cell 33:11–26.

Van de Peer Y, Mizrachi E, Marchal K. 2017. The evolutionary significance of polyploidy. Nat. Rev. Genet. 18:411–424.

Verslues PE, Agarwal M, Katiyar-Agarwal S, Zhu J, Zhu J-K. 2006. Methods and concepts in quantifying resistance to drought, salt and freezing, abiotic stresses that affect plant water status. Plant J. 45:523–539.

Vlček J, Hämälä T, Vives Cobo C, Curran E, Šrámková G, Slotte T, Schmickl R, Yant L, Kolář F. 2025. Whole-genome duplication increases genetic diversity and load in outcrossing Arabidopsis arenosa. Proc. Natl. Acad. Sci. U. S. A. 122:e2501739122.

Vyas P, Bisht MS, Miyazawa S-I, Yano S, Noguchi K, Terashima I, Funayama-Noguchi S. 2007. Effects of polyploidy on photosynthetic properties and anatomy in leaves of Phlox drummondii. Funct. Plant Biol. 34:673–682.

Wang N, Fan X, Lin Y, Li Z, Wang Y, Zhou Y, Meng W, Peng Z, Zhang C, Ma J. 2022. Alkaline stress induces different physiological, hormonal and gene expression responses in diploid and autotetraploid rice. Int. J. Mol. Sci. 23:5561.

Wang N, Wang S, Qi F, Wang Yingkai, Lin Y, Zhou Y, Meng W, Zhang C, Wang Yunpeng, Ma J. 2022. Autotetraploidization gives rise to differential gene expression in response to saline stress in rice. Plants 11:3114.

Wang Z, Liao B-Y, Zhang J. 2010. Genomic patterns of pleiotropy and the evolution of complexity. Proc. Natl. Acad. Sci. U.S.A. 107:18034–18039.

Wendel JF. 2015. The wondrous cycles of polyploidy in plants. Am. J. Bot. 102:1753–1756.

Whitacre J, Bender A. 2010. Degeneracy: a design principle for achieving robustness and evolvability. J. Theor. Biol. 263:143–153.

Wickham H. 2016. ggplot2: Elegant Graphics for Data Analysis. Available from: https://ggplot2.tidyverse.org

Wos G, Mořkovská J, Bohutínská M, Šrámková G, Knotek A, Lučanová M, Španiel S, Marhold K, Kolář F. 2019. Role of ploidy in colonization of alpine habitats in natural populations of Arabidopsis arenosa. Ann. Bot. 124:255–268.

Wos G, Požárová D, Kolář F. 2023. Role of phenotypic and transcriptomic plasticity in alpine adaptation of Arabidopsis arenosa. Mol. Ecol. 32:5771–5784.

Wu S, Han B, Jiao Y. 2020. Genetic Contribution of Paleopolyploidy to Adaptive Evolution in Angiosperms. Mol. Plant 13:59–71.

Xu S, Guo Z, Feng X, Shao S, Yang Y, Li J, Zhong C, He Z, Shi S. 2023. Where whole-genome duplication is most beneficial: Adaptation of mangroves to a wide salinity range between land and sea. Mol. Ecol. 32:460–475.

Yang PM, Huang QC, Qin GY, Zhao SP, Zhou JG. 2014. Different drought-stress responses in photosynthesis and reactive oxygen metabolism between autotetraploid and diploid rice. Photosynthetica 52:193–202.

Yao Y, Carretero-Paulet L, Van de Peer Y. 2019. Using digital organisms to study the evolutionary consequences of whole genome duplication and polyploidy. PLoS One 14:e0220257.

Yu Z, Haberer G, Matthes M, Rattei T, Mayer KFX, Gierl A, Torres-Ruiz RA. 2010. Impact of natural genetic variation on the transcriptome of autotetraploid Arabidopsis thaliana. Proc. Natl. Acad. Sci. U. S. A. 107:17809–17814.

Zhou K, Liu B, Wang Y, Zhang X, Sun G. 2019. Evolutionary mechanism of genome duplication enhancing natural autotetraploid sea barley adaptability to drought stress. Environ. Exp. Bot. 159:44–54.

