## Supplementary fig. for "Autopolyploidization presents a transient and potential-rich window of increased transcriptional plasticity in *Arabidopsis arenosa*"

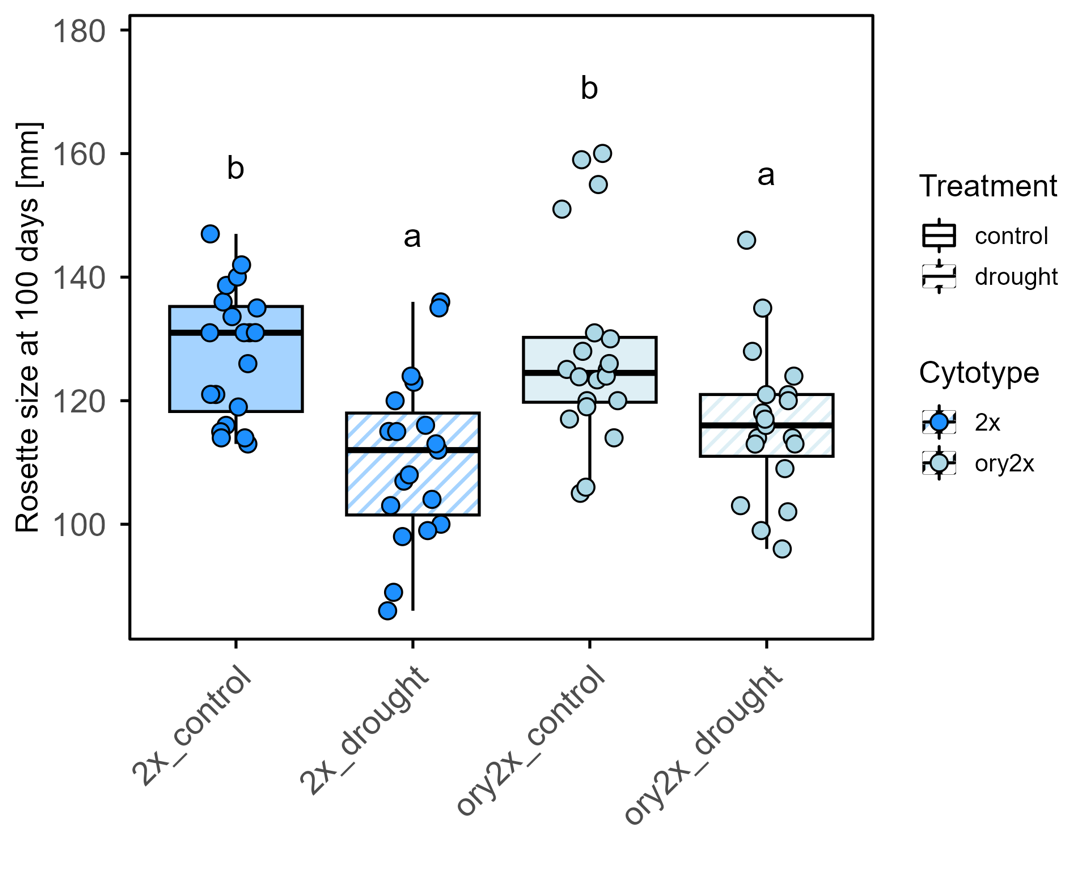

**Supplementary Figure S1. Difference in rosette diameter between F2 diploid plants and not-polyploidised discendents of oryzalin treated plants.** Rosette diameter at 100 days after sowing in natural diploid plants (2x) and oryzalin-treated diploid plants (ory2x) grown under control and drought treatments. Points represent individual plants; boxplots show the median, interquartile range, and 1.5× interquartile range. Letters above boxes denote statistically significant differences among groups based on Tukey-adjusted pairwise comparisons. Only the treatment shows an effect on rosette size.

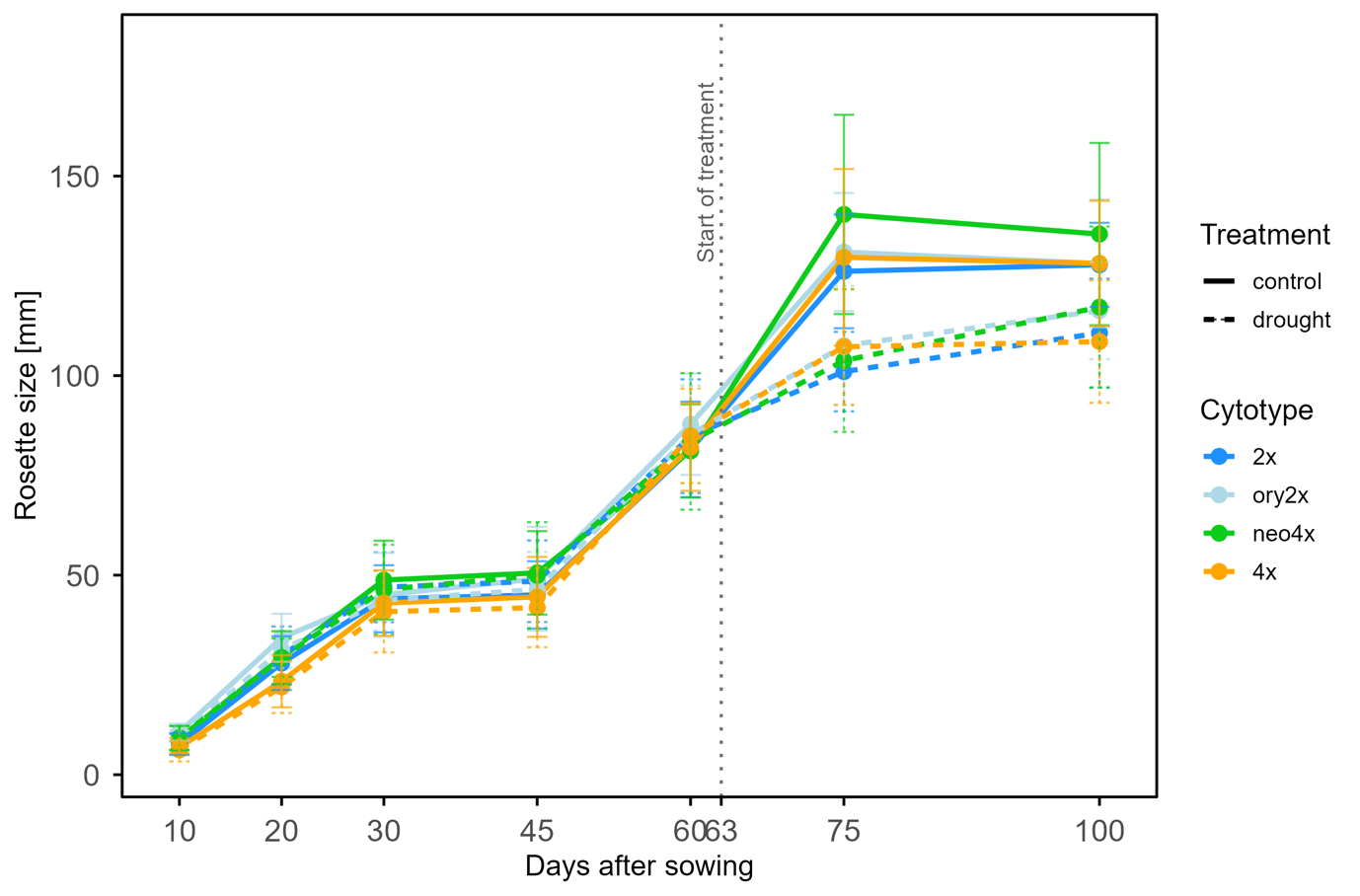

**Supplementary Figure S2. Pattern of plants vegetative growth during the time of the experiment.** Rosette diameter was measured repeatedly from 10 to 100 days after sowing across all cytotypes and treatments. Dots represent mean rosette size at each time point, and error bars indicate SD. Solid lines connect measurements from control plants, whereas dashed lines connect measurements from drought-treated plants. The vertical dotted line marks the onset of drought treatment at day 63. Growth trajectories show how cytotypes differed in early vegetative growth and in their subsequent response to drought.

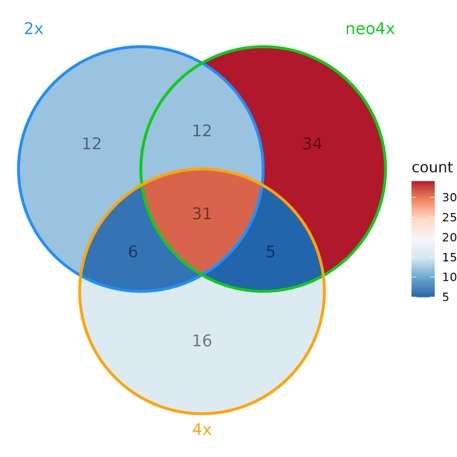

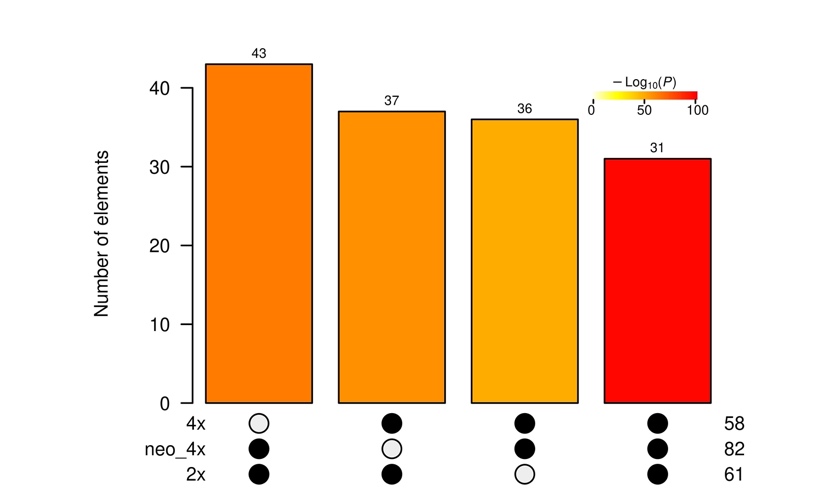

**Supplementary Figure S3. Number of significantly enriched GO-terms overlapping between ploidies DEGs.** The significance of the intersections was tested using one-sided Fisher’s exact test and it is showed on the right by the colour intensity of the bars. Each intersection shows a higher than expected by chance number of overlapping GO-terms (p < 7.429552e-50).

| **Sample ID** | **Ploidy** | **Treatment** | **Tot. reads** | **Uniquely mapped reads (%)** | **Multi-mapped reads (%)** | **Unmapped reads (%)** | **Mismatch rate per base (%)** |
| --- | --- | --- | --- | --- | --- | --- | --- |
| 8035 | neo4x | norm | 18,498,720 | 88.93 | 8.84 | 0.02 | 1.50 |
| 8129 | neo4x | norm | 15,855,967 | 90.13 | 8.65 | 0.02 | 1.52 |
| 8279 | neo4x | norm | 15,439,987 | 80.73 | 8.47 | 0.04 | 1.43 |
| 8149 | neo4x | norm | 15,413,373 | 87.55 | 8.04 | 0.03 | 1.50 |
| 8187 | neo4x | norm | 17,771,618 | 92.50 | 6.39 | 0.02 | 1.48 |
| 8219 | neo4x | norm | 15,574,733 | 88.29 | 9.52 | 0.03 | 1.52 |
| 8324 | neo4x | norm | 16,296,245 | 64.83 | 11.69 | 0.02 | 1.30 |
| 8286 | neo4x | norm | 16,400,732 | 85.05 | 7.93 | 0.03 | 1.47 |
| 8253 | neo4x | norm | 16,830,327 | 91.22 | 6.85 | 0.02 | 1.46 |
| 8055 | neo4x | norm | 18,331,527 | 80.91 | 7.46 | 0.02 | 1.42 |
| 8242 | neo4x | norm | 18,481,478 | 88.41 | 8.60 | 0.02 | 1.49 |
| 8096 | neo4x | norm | 15,325,917 | 91.02 | 6.52 | 0.03 | 1.48 |
| 8164 | neo4x | norm | 15,878,081 | 88.35 | 8.21 | 0.02 | 1.49 |
| 8092 | neo4x | norm | 19,032,397 | 89.12 | 8.88 | 0.03 | 1.49 |
| 8061 | neo4x | norm | 15,105,746 | 89.58 | 7.48 | 0.03 | 1.47 |
| 8037 | 2x | norm | 20,330,421 | 89.51 | 8.53 | 0.02 | 1.51 |
| 8077 | 2x | norm | 21,936,007 | 90.56 | 8.42 | 0.02 | 1.49 |
| 8110 | 2x | norm | 16,534,365 | 90.20 | 7.99 | 0.02 | 1.51 |
| 8255 | 2x | norm | 17,497,994 | 91.70 | 6.36 | 0.03 | 1.50 |
| 8094 | 2x | norm | 18,809,799 | 90.61 | 7.69 | 0.01 | 1.49 |
| 8186 | 2x | norm | 19,203,180 | 88.81 | 7.31 | 0.03 | 1.46 |
| 8277 | 2x | norm | 16,971,319 | 90.36 | 6.70 | 0.01 | 1.44 |
| 8246 | 2x | norm | 19,301,262 | 93.02 | 6.19 | 0.02 | 1.45 |
| 8192 | 2x | norm | 21,787,557 | 87.65 | 8.71 | 0.02 | 1.51 |
| 8172 | 2x | norm | 18,370,717 | 90.49 | 7.95 | 0.03 | 1.49 |
| 8214 | 2x | norm | 16,913,220 | 90.17 | 7.75 | 0.02 | 1.49 |
| 8287 | 2x | norm | 19,500,854 | 84.67 | 10.21 | 0.03 | 1.47 |
| 8040 | 2x | norm | 22,038,313 | 87.79 | 9.27 | 0.02 | 1.48 |
| 8328 | 2x | norm | 19,962,237 | 91.83 | 6.08 | 0.02 | 1.46 |
| 8321 | 2x | norm | 15,688,807 | 76.09 | 8.96 | 0.02 | 1.39 |
| 8278 | 4x | norm | 19,095,664 | 89.40 | 8.97 | 0.02 | 1.53 |
| 8076 | 4x | norm | 18,051,937 | 92.48 | 6.35 | 0.02 | 1.50 |
| 8044 | 4x | norm | 20,216,639 | 83.00 | 10.94 | 0.03 | 1.51 |
| 8091 | 4x | norm | 17,607,113 | 88.66 | 9.01 | 0.02 | 1.49 |
| 8121 | 4x | norm | 20,099,213 | 87.69 | 10.51 | 0.02 | 1.54 |
| 8291 | 4x | norm | 16,316,886 | 89.06 | 7.80 | 0.02 | 1.49 |
| 8176 | 4x | norm | 18,180,849 | 90.60 | 7.74 | 0.02 | 1.52 |
| 8262 | 4x | norm | 15,587,222 | 84.82 | 10.89 | 0.02 | 1.52 |
| 8213 | 4x | norm | 18,256,776 | 88.20 | 9.11 | 0.02 | 1.53 |
| 8283 | 4x | norm | 17,712,021 | 86.58 | 10.23 | 0.04 | 1.54 |
| 8197 | 4x | norm | 19,518,748 | 89.16 | 8.62 | 0.03 | 1.50 |
| 8132 | 4x | norm | 15,635,590 | 81.83 | 8.98 | 0.02 | 1.48 |
| 8248 | 4x | norm | 17,189,598 | 87.14 | 9.69 | 0.03 | 1.50 |
| 8326 | 4x | norm | 16,126,117 | 86.60 | 7.86 | 0.03 | 1.48 |
| 8238 | 4x | norm | 19,808,353 | 87.41 | 9.08 | 0.02 | 1.52 |
| 8382 | neo4x | dry | 16,544,955 | 89.91 | 7.01 | 0.03 | 1.45 |
| 8345 | neo4x | dry | 15,888,300 | 89.19 | 7.20 | 0.03 | 1.46 |
| 8479 | neo4x | dry | 21,001,851 | 90.25 | 8.36 | 0.02 | 1.49 |
| 8342 | neo4x | dry | 15,261,429 | 87.72 | 7.67 | 0.02 | 1.47 |
| 8462 | neo4x | dry | 15,262,774 | 88.45 | 7.99 | 0.02 | 1.46 |
| 8514 | neo4x | dry | 20,039,153 | 89.83 | 8.64 | 0.02 | 1.52 |
| 8541 | neo4x | dry | 22,012,277 | 90.17 | 8.56 | 0.03 | 1.47 |
| 8557 | neo4x | dry | 20,198,997 | 90.56 | 8.19 | 0.02 | 1.50 |
| 8438 | neo4x | dry | 16,592,211 | 89.13 | 7.55 | 0.02 | 1.47 |
| 8464 | neo4x | dry | 17,101,803 | 65.58 | 11.75 | 0.02 | 1.25 |
| 8331 | neo4x | dry | 16,548,693 | 89.42 | 8.95 | 0.02 | 1.52 |
| 8401 | neo4x | dry | 15,057,976 | 90.38 | 6.82 | 0.02 | 1.47 |
| 8571 | neo4x | dry | 19,409,884 | 85.95 | 9.97 | 0.03 | 1.43 |
| 8446 | neo4x | dry | 21,496,333 | 87.34 | 8.78 | 0.02 | 1.48 |
| 8412 | neo4x | dry | 19,111,621 | 87.82 | 7.47 | 0.04 | 1.47 |
| 8388 | 2x | dry | 19,118,038 | 91.23 | 6.23 | 0.01 | 1.46 |
| 8362 | 2x | dry | 18,629,287 | 91.49 | 6.41 | 0.03 | 1.44 |
| 8531 | 2x | dry | 16,879,940 | 90.00 | 6.48 | 0.01 | 1.43 |
| 8389 | 2x | dry | 19,138,363 | 88.37 | 10.19 | 0.02 | 1.51 |
| 8565 | 2x | dry | 19,032,925 | 89.25 | 8.72 | 0.01 | 1.51 |
| 8453 | 2x | dry | 20,007,956 | 87.91 | 9.77 | 0.04 | 1.50 |
| 8367 | 2x | dry | 16,841,486 | 85.92 | 7.59 | 0.02 | 1.44 |
| 8436 | 2x | dry | 19,361,427 | 88.69 | 8.95 | 0.02 | 1.50 |
| 8431 | 2x | dry | 15,675,586 | 86.26 | 9.44 | 0.03 | 1.51 |
| 8627 | 2x | dry | 18,037,261 | 89.45 | 9.04 | 0.02 | 1.53 |
| 8457 | 2x | dry | 20,116,136 | 90.45 | 7.96 | 0.04 | 1.50 |
| 8420 | 2x | dry | 15,914,067 | 87.51 | 6.97 | 0.02 | 1.44 |
| 8466 | 2x | dry | 20,044,612 | 86.11 | 11.04 | 0.07 | 1.52 |
| 8532 | 2x | dry | 18,935,154 | 81.57 | 10.16 | 0.02 | 1.46 |
| 8341 | 2x | dry | 19,253,098 | 89.12 | 8.85 | 0.02 | 1.48 |
| 8489 | 4x | dry | 16,948,441 | 89.10 | 8.94 | 0.03 | 1.54 |
| 8561 | 4x | dry | 16,728,491 | 90.96 | 7.36 | 0.01 | 1.48 |
| 8535 | 4x | dry | 16,717,342 | 89.16 | 8.78 | 0.03 | 1.51 |
| 8423 | 4x | dry | 19,620,073 | 89.12 | 9.40 | 0.02 | 1.52 |
| 8399 | 4x | dry | 19,006,407 | 89.25 | 7.57 | 0.03 | 1.47 |
| 8333 | 4x | dry | 18,988,500 | 88.89 | 9.13 | 0.02 | 1.53 |
| 8568 | 4x | dry | 17,923,276 | 85.21 | 10.58 | 0.05 | 1.53 |
| 8577 | 4x | dry | 16,809,871 | 86.90 | 10.26 | 0.02 | 1.54 |
| 8405 | 4x | dry | 19,306,615 | 90.33 | 7.86 | 0.02 | 1.50 |
| 8418 | 4x | dry | 16,226,346 | 84.51 | 8.07 | 0.03 | 1.49 |
| 8587 | 4x | dry | 20,454,821 | 89.39 | 8.94 | 0.02 | 1.50 |
| 8609 | 4x | dry | 1,194,128 | 50.03 | 30.96 | 0.21 | 1.71 |
| 8339 | 4x | dry | 17,261,289 | 88.22 | 9.40 | 0.03 | 1.50 |
| 8604 | 4x | dry | 18,887,433 | 87.05 | 10.72 | 0.03 | 1.56 |
| 8459 | 4x | dry | 15,660,256 | 86.83 | 8.11 | 0.02 | 1.50 |

**Supplementary Table S1. Summary of RNA-seq alignment and quality metrics for all samples.** For each sample, we report total number of input reads, percentage of uniquely mapped reads, percentage of multi-mapping and unmapped reads, and mismatch rate per base as obtained from STAR alignment logs. One established tetraploid sample under drought conditions (8609) showed markedly reduced mapping efficiency and elevated multi-mapped reads, consistent with low RNA quality, and was therefore excluded from downstream analyses. All remaining samples showed consistent and high-quality alignment statistics across ploidy levels and treatments.

| **Group 1** | **Group2** | **Bs. median group 1; 2.5–97.5% quantiles** | **Bs. median group 2; 2.5–97.5% quantiles** | ***W*** | ***p-value <*** |
| --- | --- | --- | --- | --- | --- |
| neo4x_dry | 2x_dry | 10,428; 6,515–14,873 | 8,552; 6,347–10,483 | 234,735 | 2.2 × 10⁻¹⁶ |
| neo4x_dry | 4x_dry | 10,428; 6,515–14,873 | 7,694; 5,684–9,389 | 868,786 | 2.2 × 10⁻¹⁶ |
| 2x_dry | 4x_dry | 8,552; 6,347–10,483 | 7,694; 5,684–9,389 | 724,010 | 2.2 × 10⁻¹⁶ |
| neo4x_norm | 2x_norm | 8,021; 5,551–10,201 | 6,365; 5,049–7,443 | 107,727 | 2.2 × 10⁻¹⁶ |
| neo4x_norm | 4x_norm | 8,021; 5,551–10,201 | 5,584; 3,766–7,399 | 941,199 | 2.2 × 10⁻¹⁶ |
| 2x_norm | 4x_norm | 6,365; 5,049–7,443 | 5,584; 3,766–7,399 | 743,379 | 2.2 × 10⁻¹⁶ |
| 2x_dry | 2x_norm | 8,552; 6,347–10,483 | 6,365; 5,049–7,443 | 955,598 | 2.2 × 10⁻¹⁶ |
| neo4x_dry | neo4x_norm | 10,428; 6,515–14,873 | 5,584; 3,766–7,399 | 818,961 | 2.2 × 10⁻¹⁶ |
| 4x_dry | 4x_norm | 7,694; 5,684–9,389 | 5,584; 3,766–7,399 | 928,564 | 2.2 × 10⁻¹⁶ |

**Supplementary Table S2.** Pairwise Wilcoxon rank-sum tests comparing transcriptomic disparity among ploidy levels and treatments.

| **GO-term ID** | **Description** | **Gene Count** | ***p-adj*** |
| --- | --- | --- | --- |
| GO:0042335 | cuticle development | 5 | 0.000838671 |
| GO:0051274 | beta-glucan biosynthetic process | 9 | 0.012994094 |
| GO:0160062 | cutin-based cuticle development | 7 | 0.012994094 |
| GO:0000271 | polysaccharide biosynthetic process | 16 | 0.012994094 |
| GO:0000910 | cytokinesis | 10 | 0.012994094 |
| GO:0030244 | cellulose biosynthetic process | 8 | 0.012994094 |
| GO:0051273 | beta-glucan metabolic process | 9 | 0.012994094 |
| GO:0030243 | cellulose metabolic process | 8 | 0.01976332 |
| GO:0016051 | carbohydrate biosynthetic process | 19 | 0.022116012 |
| GO:0009833 | plant-type primary cell wall biogenesis | 5 | 0.022116012 |
| GO:0010143 | cutin biosynthetic process | 5 | 0.022116012 |
| GO:0046467 | membrane lipid biosynthetic process | 8 | 0.028303472 |
| GO:0006643 | membrane lipid metabolic process | 9 | 0.028303472 |
| GO:2001006 | regulation of cellulose biosynthetic process | 4 | 0.039978114 |
| GO:0032950 | regulation of beta-glucan metabolic process | 4 | 0.042818516 |
| GO:0032951 | regulation of beta-glucan biosynthetic process | 4 | 0.042818516 |
| GO:0051301 | cell division | 15 | 0.042818516 |
| GO:0005976 | polysaccharide metabolic process | 21 | 0.046746897 |

**Supplementary Table S3. Gene Ontology (GO) Biological Process enrichment analysis of genes belonging to the subset of 12 neo4x modules preserved in the established tetraploid (4x) network but not in the diploid (2x) network.** GO enrichment was performed using the R package clusterProfiler v.4.14.6 on genes contained within the 12 neo4x modules. These modules comprised a total of 713 *A. arenosa* genes, of which 580 could be assigned to *Arabidopsis thaliana* orthologs and were therefore retained for downstream enrichment analyses. The table reports significantly enriched GO Biological Process terms, including the GO-term identifier, term description, number of associated genes, and Benjamini–Hochberg adjusted p-value (*p-adj*).
